# Evaluating Regulatory Module Function within Mitochondrial Pyruvate Dehydrogenase Complex

**DOI:** 10.1101/2025.11.24.690278

**Authors:** Caroline Keller, Celine Caseys, Daniel J. Kliebenstein

## Abstract

Regulatory networks coordinate metabolism to control how plants adapt to biotic and abiotic stresses. This coordination can align transcriptional shifts across metabolic pathways using cis-regulatory elements shared across the enzyme genes within these pathways. While the role of transcription factors (TFs) in controlling this process across pathways is well known, less is known regarding the role of shared cis-regulatory elements across the genes in a pathway. Sharing cis-regulatory elements across the genes in an enzyme complex or pathway, can create coordinated regulation of the pathway by a TF. However, it is unclear if all the genes in a pathway or enzyme complex need to be fully coordinated for maximal function. For example, if one gene in an enzyme complex loses a cis-regulatory element, it may not alter the function of the enzyme complexes function if post-transcriptional or compensatory transcriptional changes are sufficient to balance the complex. To test how cis-modular membership shapes the function of an enzyme complex, we used CRISPR/Cas9 to abolish a common cis-regulatory element across the promoters of nine genes required for the mitochondrial pyruvate dehydrogenase complex (mtPDC). This complex is composed of three apoenzymes and is a central hub coordinating carbon flow between glycolysis and the tricarboxylic acid (TCA) cycle. Different combinations of these cis-element mutations were tested across the genes in the complex in *Arabidopsis thaliana* and the created genotypes were phenotyped for altered enzyme function using digital growth analysis, disease assays, metabolomics, and transcriptomics. This analysis revealed that mutating cis-element motifs of genes in this enzyme complex produced distinct phenotypes, displaying promoter-specific buffering and modularity.

## Introduction

Modular coordination is a key concept of systems biology and regulatory network theory explaining the organization of different interacting networks (Ihmels et al., 2002; Melo et al., 2017). Networks function modularly with groups of genes regulated together as a regulon. In metabolism, a regulon is typically associated with specific biochemical pathways that create coordinated changes in metabolism. This model of metabolic regulation is well established in unicellular organisms where one or a few regulatory transcription factors (TFs) control specific processes. For example, *Saccharomyces cerevisiae* and *Escherichia coli* have TFs discretized into separate modules to control distinct cellular processes (Ihmels et al., 2002, 2004). In yeast, these processes often have main regulatory TFs like the Hap2/3/4/5 complex and Rtg1/2/3 system that regulate the tricarboxylic acid (TCA) cycle (Butow, 1999; Schu, 2003). Furthermore, metabolic pathways in these unicellular organisms are often organized into modules governed by central regulatory switches involving one or a few TFs (Ihmels et al., 2002, 2004; Tang et al., 2021). However, it is not clear if such pathway specific regulatory models apply to multicellular organisms. It is possible that metabolic regulation in multicellular organisms follows a different or more complicated regulatory model reflecting the need to both partition and coordinate across cell types and development.

Similar to pathway-level coordination in unicellular organisms, plant TFs play a central role in controlling metabolism. Plant TFs interact with enzyme encoding gene promoters to shape enzymatic outputs and optimize fitness, growth, and development (Gaudinier et al., 2015). However, in contrast to unicellular models, TFs regulating metabolism typically bind promoters distributed across multiple metabolic pathways. This allows plants to coordinate the regulation of different metabolic pathways in response to stress in specific cellular compartments like coupling regulation of the Pyruvate Dehydrogenase Complex (PDC) with downstream pathways that use the PDC outputs. In parallel to TFs binding multiple pathways, individual metabolic pathways are frequently bound by and controlled by a wide diversity of TFs (Tang et al., 2021). Recent work extends the complexity of plant metabolic regulation by TFs by showing that key TFs can alter metabolic pathways without influencing the expression of all the genes in the pathway. For example, mutations in key regulatory TFs for the aliphatic glucosinolate (GSL) pathway only altered expression in half the transcripts in the pathway (B. Li et al., 2014). The expression of the other genes showed that there must be additional TFs controlling their expression and likely creating overlapping modules within the pathway (B. Li et al., 2014; Wisecaver et al., 2017). As such, TF and enzyme promoter interactions in *A. thaliana* create an intricate web of regulation that likely spreads across the metabolic network to reshape the whole system and allow for rapid interconversion of required metabolites in response to diverse environmental stimuli (Tang et al., 2021). The overlap of TF effects across pathways and incomplete effects within individual pathways complicates efforts to understand metabolic pathway regulation solely via TF mutants, as these regulators can trigger complex pleiotropic effects (Hoang et al., 2017; Lan et al., 2012; Rodríguez-Leal et al., 2017; Tang et al., 2021; Yang, 2019). However, the TF centric analysis complicates the ability to test how individual cis-regulatory elements across a metabolic pathway and associated TFs may shape plant metabolism.

In addition to transcriptional regulation, metabolic systems are often controlled post-transcriptionally through the assembly and maintenance of multi-enzyme complexes, which ensure proper complex stoichiometry and coordinated catalytic activity. One example is the multi-component PDC that contains three enzymatic subunits whose coordinated activity is essential for cellular energy production. The PDC enzyme complex is central to carbon metabolism and catalyzes the oxidative decarboxylation of pyruvate from glycolysis to produce acetyl-CoA and NADH which feed into the TCA cycle. This complex consists of the apoenzymes pyruvate dehydrogenase (E1 subunit), dihydrolipoyl transacetylase (E2 subunit) and dihydrolipoyl dehydrogenase (E3 subunit), which require association of their cofactors to complete the activated holoenzyme. Combined post-transcriptional and transcriptional effects raise the question whether all the subunit encoding genes in a holoenzyme must be equally co-expressed or whether some genes can be transcriptionally disconnected from a specific co-expression module without any consequence to enzyme function and plant phenotype (Tang et al., 2021). Residual expression from other modules and protein stoichiometry control within a holoenzyme may compensate for the mutation of a single motif within the TF module. This suggests that individual cis-element mutations could have minimal phenotypic effects, whereas mutations in multiple gene promoters may be required to produce measurable outcomes. Thus, we wanted to test how sensitive an enzymatic complex might be to mutations in cis-elements across the genes encoding the subunits. By stacking mutations, we are working to understand how effects move from individual gene to transcriptome and ultimately to more downstream phenotypes and test if the genes within this enzyme complex must be coordinated by TFs or if there is robustness within this system when some genes are dropped out of a TF regulatory module without any consequence to overall fitness.

To test how regulon membership coordinates holoenzyme function, we deleted a stress-related regulatory TF cis-element motif from the promoter regions of PDC genes in *A. thaliana* (Čermák et al., 2017). This involved creating targeted mutations of specific DNA sequences across enzyme-encoding genes to generate a mutation matrix. TF membership within other regulatory modules controlling the PDC genes remains intact, since the TF can still bind to non-mutated promoters. Eliminating an individual gene’s response to stress allows us to quantify the effect of removing individual components of the TF module in this enzyme complex and evaluate the importance of each member in the module. as *A. thaliana* is a good model system to study modularity in metabolism due to the public availability of its full genomic sequence, TF maps across the genome, transcriptional networks, protein–protein interactions, pathogen interactions, and metabolic assays (Meyer et al., 2007; Obayashi et al., 2007; Palaniswamy et al., 2006; Yilmaz et al., 2011).

We test if PDC holoenzyme subunits must be co-expressed to maintain phenotypic stability and metabolic coordination when individual and combined stress-related motifs are ablated. This approach tests the hypothesis that partial co-regulation of enzyme complexes by TF modules is sufficient to sustain primary metabolism. This research provides information on how removing a specific cis-regulatory module alters the plant’s phenotype under stress, shapes genotype-to-phenotype relationships and enables the engineering of plant metabolism through modular design.

## Materials and Methods

### Locating Modules and the Cis-Element Motifs

To identify regulatory modules to potentially target in *A. thaliana*, we combined yeast one-hybrid Y1H (Tang, Li, Zhou, Bolt, Li, Cruz, Gaudinier, Ngo, Clark-Wiest, et al. 2021) and DAP-seq datasets (O’Malley et al., 2016) describing global promoter-TF interactions. For each gene’s promoter, a matrix of 1,129 high-scoring DAP-seq motif occurrences (1 kb upstream of transcription start site) were scanned with FIMO in MEME Suite 5.5.7 (Grant, Bailey, and Noble 2011), and TF binding motifs were selected based on significance (p < 0.0001), presence across the mitochondrial pyruvate dehydrogenase complex (mtPDC) promoters, overlap with Y1H interactions, and low copy number per promoter. Promoter sequences for nine mtPDC subunits (pAT1G01090, pAT1G48030, pAT1G54220, pAT1G59900, pAT3G13930, pAT3G17240, pAT3G52200, pAT5G50850, pAT1G24180) were obtained from TAIR (http://www.arabidopsis.org) and NCBI (Sayers et al., 2024) using the TAIR10 genome assembly (Lamesch et al., 2012). The mtPDC complex was chosen as a model because it contains a complete set of nuclear encoded PDC subunits, linking energy metabolism to stress-responsive adaptation. For motif selection, stress-related TFs were prioritized to focus on cis-element mutations that should have more conditional effects, and we avoided developmentally associated TF motifs (Toufighi et al., 2005). Motifs were further filtered with the Multiple List Comparator (Baschal et al., 2020) to locate low copy number across as many promoters as possible for CRISPR/Cas9 targeting. This led us to focus on motifs connected with ATHB34 (AT3G28920), a stress-responsive TF whose mutants have PDC associated phenotypes (Tang et al., 2021). There were 13 occurrences of the known ATHB binding motif distributed across 8 of 9 mtPDC promoters. The AT1G24180 promoter had no occurrence of the ATHB binding motif within the DAPseq nor the Y1H TF-DNA interaction network of transcriptional regulation of *A. thaliana* primary and specialized metabolism (Tang, Li, Zhou, Bolt, Li, Cruz, Gaudinier, Ngo, Clark-Wiest, et al., 2021).

### CRISPR/CAS9 Design and Assembly of Constructs

CRISPR/Cas9 multiplexed single guide RNAs (sgRNAs) (Stuttmann et al., 2021) were designed to independently target the ATHB34 TF binding elements in each of the mtPDC promoters to generate mutants with ATHB motifs in each genes promoter. Each binding site motif across all PDC genes were targeted to eliminate the TF binding element. CRISPR-P V2 (Liu et al., 2017) was used to search for sgRNAs by inputting each of the target mtPDC promoter sequences, a NGG PAM (spCas9 from Streptococcus pyogenes: 5’-NGG-3’), a U6 snoRNA promoter, the preset RNA scaffold (GUUUUAGAGCUAGAAAUAGCAAGUUAAAAUAAGGCUAGUCCGUUAUCAACUUGAAAAAGUGGCACCGAGUCGGUGCUUUU), and a sgRNA sequence length of 20bp were used to design each construct. The cloning overhangs (ATTG-GTTT) on the sgRNA correspond to the U6-26 promoter fragment from *A. thaliana* (*at*U6) Dicot Genome Editing (pDGE) vectors (Ordon et al., 2017). SpCas9 NGG PAM sites were identified using CRISPR-P v2.0 (Liu et al., 2017) and sgRNAs predicted to disrupt the ATTA motif bound by ATHB34 and located in all the mtPDC promoters were selected. All sgRNAs were compared against the genome to identify sgRNAs with a low “off-score” i.e. sgRNAs mainly binding the target with no likely second site targets. The cutting frequency determination (CDF) score was calculated as described by (Doench et al., 2016) to predict the effects on any off-target sites, and the sgRNAs were filtered for a high CFD to promote better on-target activity. Additionally, all potential off-target binding sites were analyzed with NCBI’s BLAST (Altschul et al., 1990) to assess their potential effects.

The oligos containing the sgRNA sequences targeting each motif were hybridized and loaded into sgRNA shuttle vectors to create modular stacks and combinations of various sgRNAs (Stuttmann et al., 2021). The recipient vector was pDGE651 i.e. FAST_hpt_pRPS5a:Cas9_ccdB with hygromycin (hpt), spectinomycin (spec), chloramphenicol (cm), and FAST as selection markers (Stuttmann et al., 2021). Twelve different constructs with different combinations of gRNAs were created with up to 13 gRNAs total in one construct.

### Preparation and verification of plasmid DNA

Plasmids were purified using the QIAprep Spin Miniprep Kit (QIAGEN 27106), restriction digested as per the BbSI, BsaI NEB protocol and (Stuttmann et al., 2021). Electrophoresis of DNA samples was performed on 50 mL or 150 mL 0.8–2.2 % w/v agarose gels (Sigma-Aldrich A9539), DNA concentration was taken with a Thermo Scientific NanoDrop Spectrophotometer or Qubit Fluorometer. Each gRNA was cloned and stacked into the final vector, promoter constructs were verified by Sanger sequencing with Azenta Life Sciences (formerly Genewiz) using the primers as specified by (Stuttmann et al., 2021). The initial vectors were transformed in MACH1 (Invitrogen, C8620-03) competent cells as detailed by Invitrogen’s competent cell transformation protocol instructions (Invitrogen, 2006).

Validated constructs were transformed into *Agrobacterium tumefaciens* GV3101 (pMP90) (Logemann et al., 2006). A volume of 50uL of chemically competent cells was incubated with 1uL of ∼50ng plasmid on ice for 30 minutes, heat shocked for 30 seconds at 37°C and incubated on ice for another 5 minutes. For recovery, 250uL of LB media was added, and cells were incubated for 2 hours at 28°C. For selection,100uL was plated on LB agar plates + gentamicin and spectinomycin antibiotics and incubated at 28°C for 2 days.

### *A. thaliana* Transformation

*A. thaliana* Col-0 plants were grown in a 4×9 tray in long day conditions (14h light/10h night) at 20°C 60% humidity. The first flower buds were cut to promote further flowering. After 5 weeks, *A. thaliana* Col-0 flowers were floral dipped. A 5mL starter culture of the CRISPR plasmid transformed in GV3101 was grown in LB media overnight at 28°C. 1:100 starter culture was used to inoculate 500mL of LB + antibiotics and grown for ∼15-17 hours at 28°C. The agrobacterium cells were pelleted and resuspended in 10mL of 5% sucrose, 500mL water and 0.0005% silwet L77 solution. Flowers were dipped into this solution and dried on their side overnight in the dark and grown in the growth chamber until plants started producing siliques. Transformed seeds were selected by presence of red fluorescence protein (RFP) (FAST marker) in the seed. DNA from T1-T3 plants was extracted as described by (Sharma et al., 2023) and mutations were confirmed with Amplicon Sequencing as described by (Campbell et al., 2015). Plants containing as many combinations as possible of the correct mutations were moved forward each generation until homozygous mutations were located.

### *A. thaliana* Growth for all Experiments

The plants were grown in Sunshine Mix #1 produced by Sun Gro Horticulture and fertilized with N-P-K macronutrient ratio of 2:1:2, containing no boron. Controlled environmental chamber conditions were set to short day conditions (10h day/14h night at 20 °C at 100-120 mE light intensity). The 30 genotypes were grown in 4×9 trays following a randomized block design for each experiment and replicated 6-fold. As such, each block consisted of all 30 genotypes randomized across five planting trays, plus 6 genotypes randomized in a block distributed across all the trays. These plants were used to measure the effects of the cis-mutations associated with each genotype on rosette growth, production of phytochemical defenses (camalexin and glucosinolate content), transcriptome changes, and susceptibility to *Botrytis cinerea*, a fungal pathogen. All experiments were repeated 3 times independently and the same seed stock was used.

### Rosette Growth Experiment

To assess the potential impacts of the cis-element disruption in the mtPDC complex on plant growth, we measured rosette growth across all 30 genotypes. Starting one week after planting, plants were photographed every other day until their rosette leaves started overlapping. The images were used to quantify rosette size with ImageJ (Schneider et al., 2012) following the BTI ImageJ measurement protocol (BTI Curriculum Development Projects in Plant Biology, 2015).

To assess the potential impacts of the cis-element disruption in the mtPDC complex on plant growth, we measured rosette growth across all 30 genotypes. Starting one week after planting, plants were photographed every other day until their rosette leaves started overlapping. The images were used to quantify rosette size with ImageJ (Schneider et al., 2012) following the BTI ImageJ measurement protocol (BTI Curriculum Development Projects in Plant Biology, 2015). For each plant, the total leaf area [cm^2^] was tracked across time. To estimate the effect of cis-mutations, the leaf area was modeled according to the genotype and number of days since planting (Days), while correcting for experimental error.

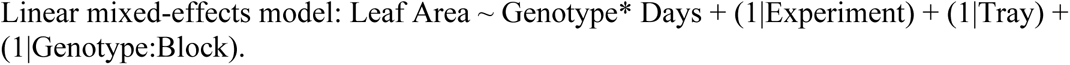

Experimental replication (1-3), trays in which the plants were grown, and the block effect were considered as random effects. The interaction term genotype × days assessed if genotypes differed in growth over time (Supplemental Table 2 and Supplemental Table 3). Given that the experimental design controlled for random factors, adjusted means were estimated using EMMeans with Satterthwaite approximation (Supplemental Table 4). To assess the difference to Col-0 (wild-type) and each mutant genotype, the pairwise contrasts were filtered for statistical significance p < 0.05. of freedom and the pairwise contrasts were FDR corrected (Supplemental Table 3).

### Susceptibility to *B. cinerea* tested by Detached Leaf Assay (DLA)

To assess the potential impacts of the cis-element disruption in the mtPDC complex on resistance to *B. cinerea*, plants were grown in short day conditions in a randomized complete block design as described above and infected with six diverse *B. cinerea* isolates as described by (Caseys et al. 2021; Zhang et al. 2017). Detached leaves from eight weeks old plants in the vegetative phase were infected with six diverse *B. cinerea* isolates named 1.03.18, 1.04.04, 1.04.17, Molly, Kern B2, and Triple7 (Caseys et al. 2021; Zhang et al. 2017). These *B. cinerea* isolates were chosen from the lab strain collection (Caseys et al., 2021) to cover a range of low to high Botrydial pathway gene expression, a fungal phytotoxin (Zhang et al. 2017). For each genotype, seven fully mature developmentally matched leaves were detached from a single plant and infected with a single drop of a *B. cinerea* isolate or a grape juice control. The infections were replicated 6-fold across experimental trays in a randomized complete block design, resulting in 1260 observations. Lesions were imaged at 72 and 96 hours post infection (HPI). The lesion area for each leaf was measured with R-scripts (Fordyce et al, 2018). It can happen that by chance, inoculations fail for unknown reasons (Caseys et al. 2021). These failed-lesions were cleaned and filtered with a threshold of mm^2^<35 corresponding to the average size of drops containing only grape juice (controls). We estimated the contributions of the interaction of *B. cinerea* isolates with plant genotypes, and experimental factors to the lesion area with the linear mixed model:

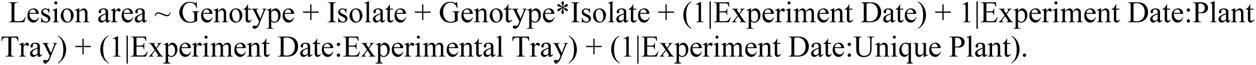

In this model, the three independent experiments (“Experiment Date”), trays in which the plants were grown (Plant tray), the experimental tray in which the detached leaf was infected (“Experimental Tray”), and the individual plants from which the leaves were detached (Unique plant) were considered as nested random effects. This model was run independently on the lesions measured at 72 hpi and 96 hpi (Supplemental Table 2). For each genotype, between 57 and 118 lesion measurements were obtained across two time points (72 hpi and 96 hpi). Unequal replication resulted from variation in infection and growth success and was accounted for in linear mixed-effects models. Post-hoc (t-test) were run within the model to compare the difference of each mutant genotype versus the wild-type genotype (Supplemental Table 3). The margin means were obtained using EMMeans (Supplemental Table 4). The EMMeans from 72HPI and 96HPI were combined into a single matrix, and Principal Component Analysis (PCA) was performed to visualize the distribution of lesion effects across the genotypes.

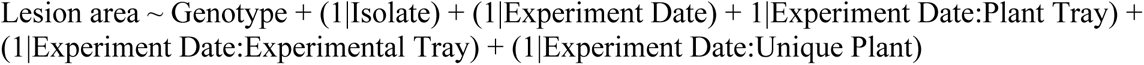

This model was used once the ANOVA revealed that there was no significant Genotype*Isolate interaction, and both Genotype and Isolate were significant. This revealed which genotypes were significantly different from the wild-type (Figure 2).

### GSL metabolite analysis

To assess the potential impacts of the cis-element mutations in the mtPDC complex on defense metabolism, GSL and camalexin accumulation was measured using leaves from the above experiments. For a developmental assay, tissues from the inner, middle, and outer whorls were taken from ∼8 week old plants for GSL metabolite analysis to measure if promoter mutant genotype and rosette position influenced GSL accumulation (Brown et al., 2003; Hunziker et al., 2021; Púčiková et al., 2023). Extra plants were grown in addition to this experiment and individual leaves were removed from these plants to be infected by six *B. cinerea* isolates (1.03.18, Molly, Triple7, 1.04.04, Kern B2, and 1.04.17) (Brown et al., 2003; Hunziker et al., 2021; Púčiková et al., 2023). Individual detached leaves were infected with *B. cinerea* isolates (1.03.18, Molly, Triple7, 1.04.04, Kern B2, and 1.04.17) for 48 hours and collected for metabolite profiling on an HPLC-DAD. Camalexin and GSLs were measured as previously described (Kliebenstein et al., 2005; Kliebenstein, Gershenzon, et al., 2001; Kliebenstein, Kroymann, et al., 2001; Kliebenstein, Lambrix, et al., 2001). Briefly one leaf (∼40mg) was added into 400uL 90% MeOH, it was crushed and filtered. Then the leaf was homogenized and extracted as described by (Katz et al., 2021). The leaf area was digitally measured for future use to standardize amounts across samples. The resulting extract was passed over the Sephadex DEAE A25 (Sigma-Aldrich) to retain the glucosinolates. This initial flowthrough of 90% MeOH was retained and analyzed for camalexin (only plants infected with *B. cinerea*). Camalexin was detected with a fluorescence detector at emission 318 nm/excitation 385 nm. The remaining sample represented GSLs bound to Sephadex and was incubated overnight with sulfatase to release the GSL, and the flowthrough was analyzed for GSL content by HPLC-DAD as described by (Katz et al. 2021) and (Caseys et al., 2024). All compounds were identified and quantified using known standards (Jensen et al., 2015) and the GSL and camalexin amounts across samples were standardized to units per leaf area as previously conducted (Rowe et al., 2010).

This allowed the quantification of numerous GSL traits including the aliphatic GSLs: 3-methylthiopropyl (3MT), 3-methylsulfinylpropyl (3MSO), 4-methylsulfinylbutyl (4MSO), 5-(methylsulfinyl)pentyl (5MSO), 7-methylsulfinylheptyl (7MSO), 4-methylthiobutyl (4MT), 8-methylsulfinyloctyl (8MSO), and the indolic GSLs: indol-3-ylmethyl (I3M) and 4-methoxy-3-indolylmethyl (4MOI3M). As additional traits, the sum of all GSLs in sample, aliphatic GSLs, indolic GSLs, C3 GSLs, C4 GSLs, short chain (3C + 4C), long chain (7C + 8C), the ratio of 4MT/4MSO+4MT that measures the oxidation state, the C7:C8 Ratio, the ratio of 4MOI3M/I3M+4MOI3M that relates to jasmonic acid signaling, and the short to long chain ratio and camalexin (for the infected leaves) were calculated.

For all GSLs, linear mixed models were fit using the lmer() function from the lme4 package, and model significance was assessed via type III ANOVA using Satterthwaite approximation for degrees of freedom. Percent variance explained (PVE) was calculated for each fixed effect. Each model is detailed as follows:

Using the glucosinolate data obtained from different leaves within the rosette we ran the following model:

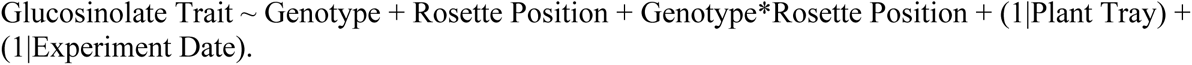

Rosette positions correspond to the inner, middle, and outer parts of the rosette. This model used the genotype x rosette position terms as fixed effects, and post-hoc comparisons (t-test) of each mutant genotype versus wild-type were conducted independently within each rosette position and EMMeans obtained (Supplemental Tables 2-4).

Using the stress-induced glucosinolate data we ran the following model: Glucosinolate Trait ∼ Genotype + Isolate + genotype*isolate + (1 | Plant Tray). This model was performed on metabolites measured from leaves infected with the six *B. cinerea* isolates or the mock grape juice control after 48 hours. This model used the genotype x isolate term as fixed effects and post-hoc comparisons (t-test) of each mutant genotype versus wild-type were conducted independently within each isolate and EMMeans obtained (Supplemental Tables 2-4).

To measure the effect of the *B. cinerea* infection on phytochemical defense, we ran the following model: Glucosinolate Trait ∼ Genotype + Treatment + genotype*treatment + (1 | Plant Tray). This model tested for genotype effects on metabolites in response to infection with *B. cinerea* (yes infected with *B. cinerea* or no “treated” with a mock grape juice control). This model used the genotype x treatment term as fixed effects and post-hoc comparisons (t-test) of each mutant genotype versus wild-type were conducted independently within each treatment and EMMeans obtained (Supplemental Tables 2-4).

### Transcriptomics

Leaf tissue from the same plants grown for the detached leaf assay *B. cinerea* experiments were collected and immediately frozen in liquid nitrogen for RNA sequencing each time the experiment was performed (one uninfected leaf from each genotype in every tray over three replicated experiments). A total of 180 mRNA libraries were prepared for paired-end sequencing representing 3x replication across each of the 30 genotypes. Libraries were prepared as previously described by (Kumar et al., 2012) with minor modifications from (Zhang et al., 2017, 2019) using NEXTex-96-RNA-Seq Barcodes. The pre-pooled libraries were size selected for ∼300bp, pooled in 90-sample batches, and sequenced at UC Davis Sequencing center (DNA Technologies Core, Davis, CA) and Expression Analysis Core Facility Genome Center. We used Aviti PE 150 Sequencing with 8bp indexing using standard Illumina Primers. Raw reads were quality controlled with MultiQC v1.15 and low-quality samples were removed. Specifically, each library was required to contain a minimum of a million reads to be included for analyses. Reads were aligned to the *A. thaliana* TAIR 10 reference genomes with HISAT2, and Rsubread to generate read count files. Normalization of the gene expression data was performed with EdgeR filtering lowly expressed genes with a count-per-million (CPM) value of > 0.5 in at least two samples. Gene counts passing this filter were further normalized with calcNormFactors() using the Trimmed Mean of M-values (TMM). The R/Bioconductor software package Limma (Ritchie et al., 2015) was used to analyze gene expression. Two different models were utilized to analyze gene expression based on either shared promoter motif mutation or specific mutant genotype. The first model (expression ∼ motif / genotype) nested each genotype within the targeted motif in each promoter as either wild-type or mutant. This model was used to generate the weighted average difference in the average expression for each mutated motif versus the wild type while controlling for background genotype. The second model (expression ∼ genotype) compared the overall difference between each genotype and the wild-type control, utilizing G30 as the wild-type baseline. This generated the differential expression of each genotype vs. the wild type. Both linear models were fit with lmfit, eBays was used, and DE genes were extracted with topTable within the Limma package. The linear modeling was performed within Limma and pairwise genotype comparisons (t-test) were generated with makeContrasts i.e. to located differentially expressed genes in each genotype relative to G30 (the same was performed with the motif model).

The pairwise contrasts for the genotype and motif model generated via Limma were used for GO enrichment. This was performed by filtering genes with adjusted p-values < 0.05, mapped *A. thaliana* gene identifiers (TAIR) to Entrez IDs and conducted biological process (BP) enrichment using the clusterProfiler package in R (Supplemental Table 5 and 6). The significant GO terms located for each genotype contrast to G30 were included in Figure 8.

### Statistical Analysis

All statistical analysis and visualization were performed using R Statistical Software (v4.2.2; R Core Team 2022). Lesion area, metabolite concentrations, growth, and gene expression were each modeled independently for each experiment using linear mixed models with the lme4 package. The EMMeans and contrast() functions were also utilized for data analysis. The prcomp function within base R was utilized to perform PCA on each of the datasets to visualize the data structure.

## Results

### Design and Assembly of Constructs Targeting mtPDC ATHB Cis-Regulatory Element Binding Sites

To begin assessing the role of cis-regulatory element coordination in plants, we focused on the mitochondrial pyruvate dehydrogenase complex in *A. thaliana*. The mtPDC enzyme complex is encoded by nine genes, some of which also alternatively localize in the cytosol and plastid (Duncan et al., 2011; Song et al., 2024). To identify candidate TFs, we integrated Y1H and DAP-seq data to detect shared motifs across the promoters and identify their associated TFs. This method identified two candidate TF/motif sets, one MYB and one ATHB, present at 1-2 copy numbers in most mtPDC gene promoters. Previous work has shown that ATHB34 likely regulates mtPDC activity and binds to promoters of TCA cycle genes as part of a transcription factor–TCA (TF–TCA) regulatory network. Moreover, the TCA cycle function is altered in *ATHB34* mutants (Tang et al., 2021). We targeted the ATHB binding sites across the mtPDC promoters to create homozygous motif mutations. After screening over 200 progeny across 13 different motif sites (Table 1), we identified 29 mutant lines containing various mutations in different ATHB34 binding sites across the mtPDC promoters (Table 1, Figure 1). Although all the sgRNAs had comparable predicted efficiencies, mutation frequency was uneven across motif targets. Four of the 13 motifs yielded no detectable mutations while two motifs were frequently mutated. This variability did not correlate with promoter redundancy. For example, the AT3G52200 promoter, which contains three motifs, might be expected to be more amenable to mutation due to redundancy, yet only three mutation lines were obtained. In contrast, the AT3G13930 promoter, which contains a single ATHB motif, yielded 21 independent mutations. In cases where a promoter contained multiple ATHB motifs, we observed preferential mutation of one motif over another by the same sgRNA. Altogether, mutational bias might be influenced by the gene context rather than the redundancy of the promoter motif. Using this matrix of mutations in ATHB binding motifs across mtPDC promoters, we phenotyped the 29 mutant lines across various conditions. This was done to analyze how variation in motif number and location influences promoter responsiveness under stress conditions like exposure to a pathogen and how altering this core metabolic pathway can influence diverse metabolic, growth and defense phenotypes.

**Figure 1:**
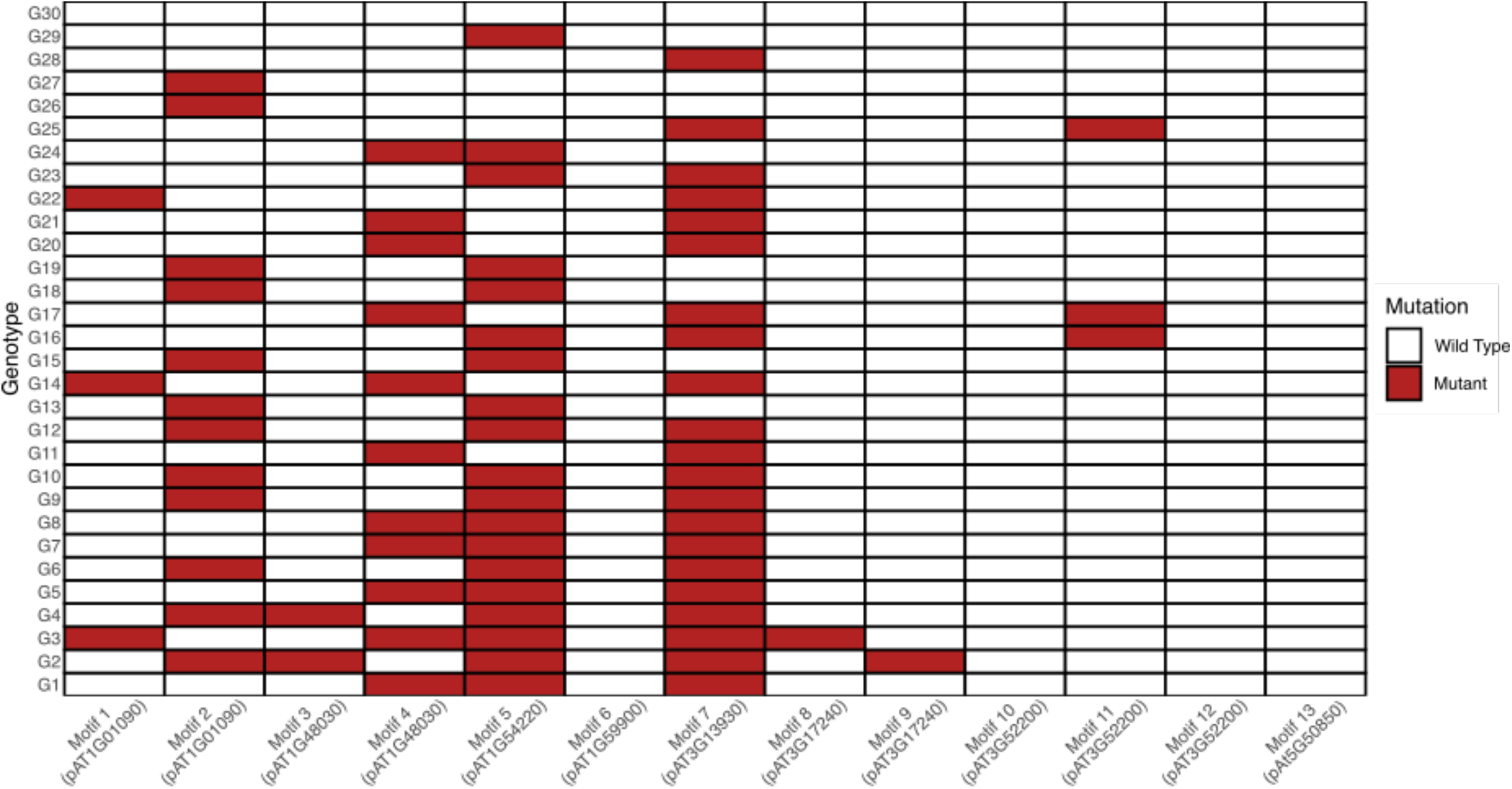
Mutation Matrix; 29 genotypes with homozygous mutations in one to five cis-elements were generated by Crispr-Cas9 editing. Successfully edited motifs are in red. Wild-type motifs for which no mutations were generated due to ineffective sgRNAs or the inability to disrupt the motif are in white.

**Figure 2:**
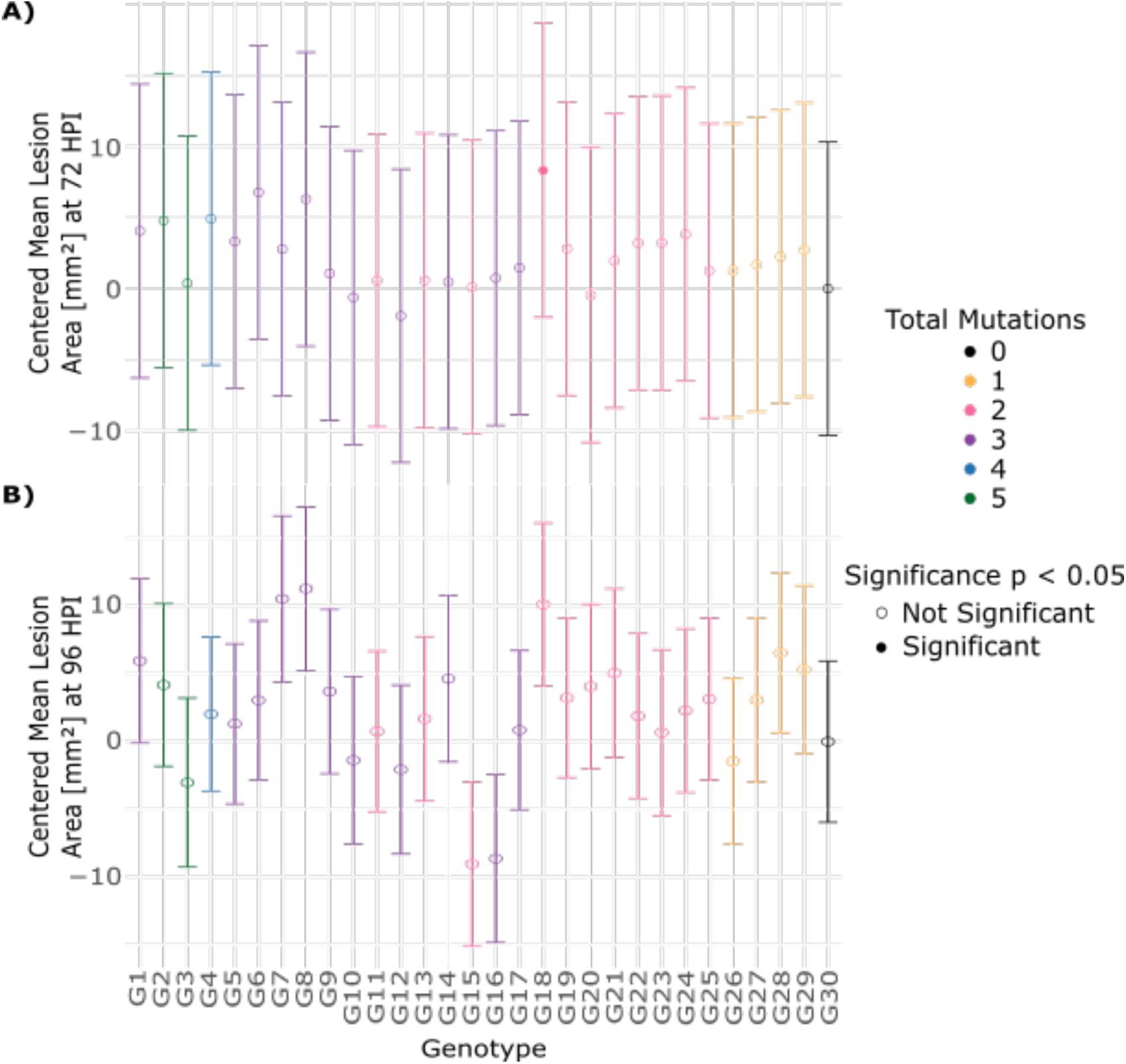
Susceptibility of the genotypes to *B. cinerea* utilizing the DLA Genotype Mean Lesion Area at both 72 HPI and 96 HPI. To visualize the effects of mutations independently of the background, the lesion area of each genotype was centered by subtraction of the mean lesion area of wild-type. As such, negative values indicate a decrease in susceptibility and positive values an increase in susceptibility compared to wild-type. Mean and +/- 1 standard error (SE) of the EMM of the lesion area at A) 72 HPI and B) 96 HPI. The solid circles indicate significant difference (FDR<0.05) to wild-type, The Satterthwaite approximation was used for degrees of freedom and the pairwise contrasts were FDR-adjusted.

**Table 1:**
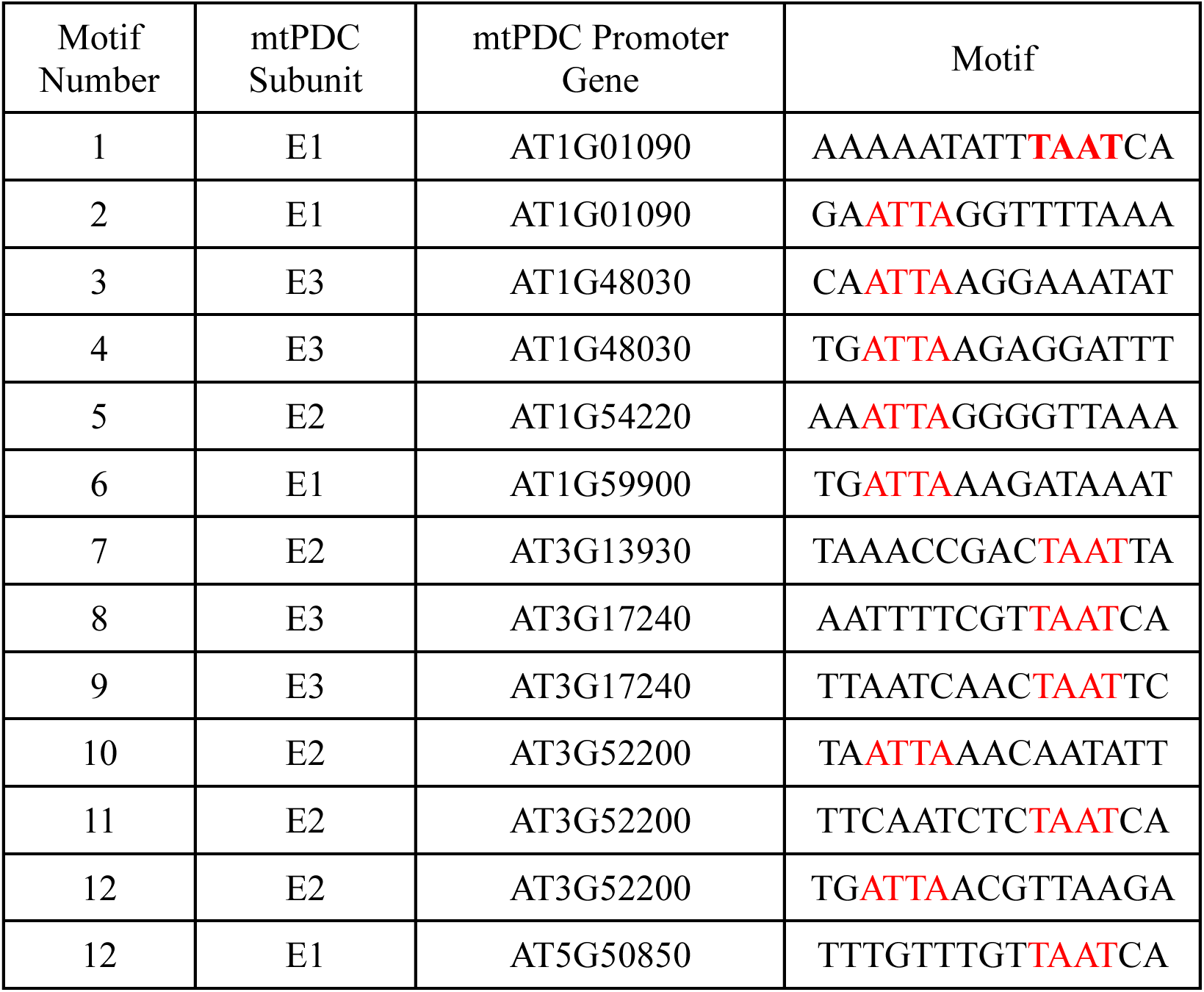
Cis-element motifs associated with eight genes encoding three subunits of the mtPDC bound by the TF ATHB34. These motifs were selected for their broad presence across mtPDC promoters and previous validation by Y1H. This table displays the motif binding sites bound by TF ATHB (in red), the mtPDC subunit each promoter gene is bound by, and the number of successful Crispr-Cas9 edits.

### The Effects of ATHB Cis-regulation of mtPDC on Rosette Growth

The mtPDC complex, a key component of the TCA cycle, plays a central role in carbon flux and general plant metabolism. Given its importance, we wanted to ascertain if cis-element mutations in mtPDC gene promoters affect plant growth (Yu et al., 2012). To test this, we measured rosette growth across all 29 mutant lines and wild-type starting at 13 days after planting (DAP) to 34 DAP. We used a linear mixed-effects model to analyze the leaf area growth across DAP for each genotype. Five cis-element mutants (G9, G10, G12, G21, and G29) showed a consistent and significant increase in growth (Figure 4), which was the most pronounced in the later stages of development. The strongest effect was observed in G10, which had an increased leaf area of 1.66 cm² at 23 DAP in leaf area (cm^2^) and 3.77 cm² at 34 DAP (p = 4.3E-13) compared to wild-type (Supplemental Table 3). These four genotypes shared mutations in three specific gene promoters (AT1G01090, AT1G54220, and AT3G13930), suggesting that altering the ATHB regulation of these three genes simultaneously leads to an almost epistatic effect on growth. This supports the idea that ATHB34-mediated regulation of mtPDC genes contributes to plant growth, and that coordinated disruption of multiple promoter elements is required to observe significant phenotypic effects. However, not all combinations of promoter mutations yielded the same outcome, highlighting the complexity of this regulatory network.

### The effects of ATHB cis-regulation of mtPDC on stress-resistance

The enzymatic role of the mtPDC complex within the TCA cycle acts as a central metabolic hub that feeds into the shikimate pathway, responsible for the production of a diversity of aromatic amino acids and phenylpropanoids contributing to plant defense. Prior studies have shown that ATHB34 TF mutants have increased resistance to *B. cinerea,* potentially linked to jasmonic acid signaling and growth vs. defense tradeoffs in immune response (X. Li et al., 2025). Given this connection, we tested if ATHB-binding motif mutations within the mtPDC gene promoters also alter plant disease resistance. We chose *B. cinerea* due to its known sensitivity to camalexin and other compounds derived from the shikimate pathway (Ferrari et al., 2007). We infected the 29 mutant lines and wild-type with six *B. cinerea* isolates that differed in their botrydial content (Zhang et al., 2017) and can be used to apply targeted biotic stress on *A. thaliana*. This was done to explore altered module membership of the mutants under stress by looking at the *B. cinerea* lesion sizes on leaves. To test if there might be Isolate dependent genotype effects, we ran a linear mixed model including a genotype x isolate interaction term. Genotype was weakly significant at 72 HPI (p = 0.035) and only lessened by 96 HPI (p = 0.981) and there was no significant genotype x isolate interaction effects at 72 HPI (p = 0.373) nor at 96 HPI (p = 0.739). We thus focused on testing if there was a genotype effect on general *B. cinerea* resistance across all the isolates by designating genotype as the fixed effect and making the isolate a random effect. At 72 HPI, the overall genotype effect was significant (p = 0.016) but was not significant at 96 HPI (p = 0.084) (Supplementary Table 2). Setting genotype as the main effect allowed us to run contrasts to see which genotypes were significantly different from the wild-type. Only G18 showed a statistically significant increase in lesion size compared to wild-type across all isolates (Figure 2, Supplementary Table 3).

To visualize the effects of the promoter motif mutants on disease resistance, we conducted principal component analysis using the EMMeans of the lesion size for each promoter motif mutant infected with each *B. cinerea* isolate. If all the promoter motif mutants had a similar quantitative positive or negative impact on disease resistance, we would expect a dominant principal component capturing most of the variation. Instead, the PCA revealed high diversity in the resistance profiles of the mutants, indicating a lack of a unifying directional effect (Supplementary Figure 1). G18 was the most distant from the wild type G30, confirming the linear mixed-effect model results. The PCA patterns revealed no association to the number of promoter motifs mutated but tracked specific combinations of mutations (Supplemental Figure 1). Thus, under the tested conditions, disruption of ATHB motifs within the mtPDC had at best subtle and differential effects on resistance to *B. cinerea*.

### Cis-element Mutations Subtly Affect GSL Accumulation Across Rosette Position

To directly assess if the cis-element mutations in mtPDC promoters alter defense metabolism, we measured GSL and camalexin accumulation. Indolic GSLs and camalexin are derived from tryptophan, a product of the shikimate pathway, while the aliphatic GSLs are derived from the methionine, a product of a distinct biosynthetic pathway. By profiling 18 different GSL metabolites, related traits like oxidation state, signaling pathways, and camalexin accumulation, we measured different quadrants of the *A. thaliana* metabolic defense grid. As previously reported, *A. thaliana* spatially partitions GSLs across the rosette to optimize defense in young leaves versus mature leaves, likely in an effort to optimize the metabolic cost of the defense (Brown et al., 2003; Hunziker et al., 2021; Púčiková et al., 2023). To model the effect of cis-element mutations across a developmental gradient, we measured the accumulation of each glucosinolate independently in leaves at different rosette positions for each genotype. This genotype x rosette position interaction (i.e., stage of development within the whorl) showed that both genotype and rosette position had significant effects on glucosinolate metabolism (Figure 3A). Rosette position explained a greater proportion of variation in aliphatic GSLs, while genotype had stronger effects on Indolic GSLs. Interestingly, no significant genotype x rosette position interactions were observed. As previously reported, the older outer leaves accumulated more long-chain GSLs and oxidized aliphatic GSLs, while the younger inner leaves accumulated higher levels of total GSLs and short-chain GSLs (4MSO and 5MSO) (Supplemental Table 4).

**Figure 3:**
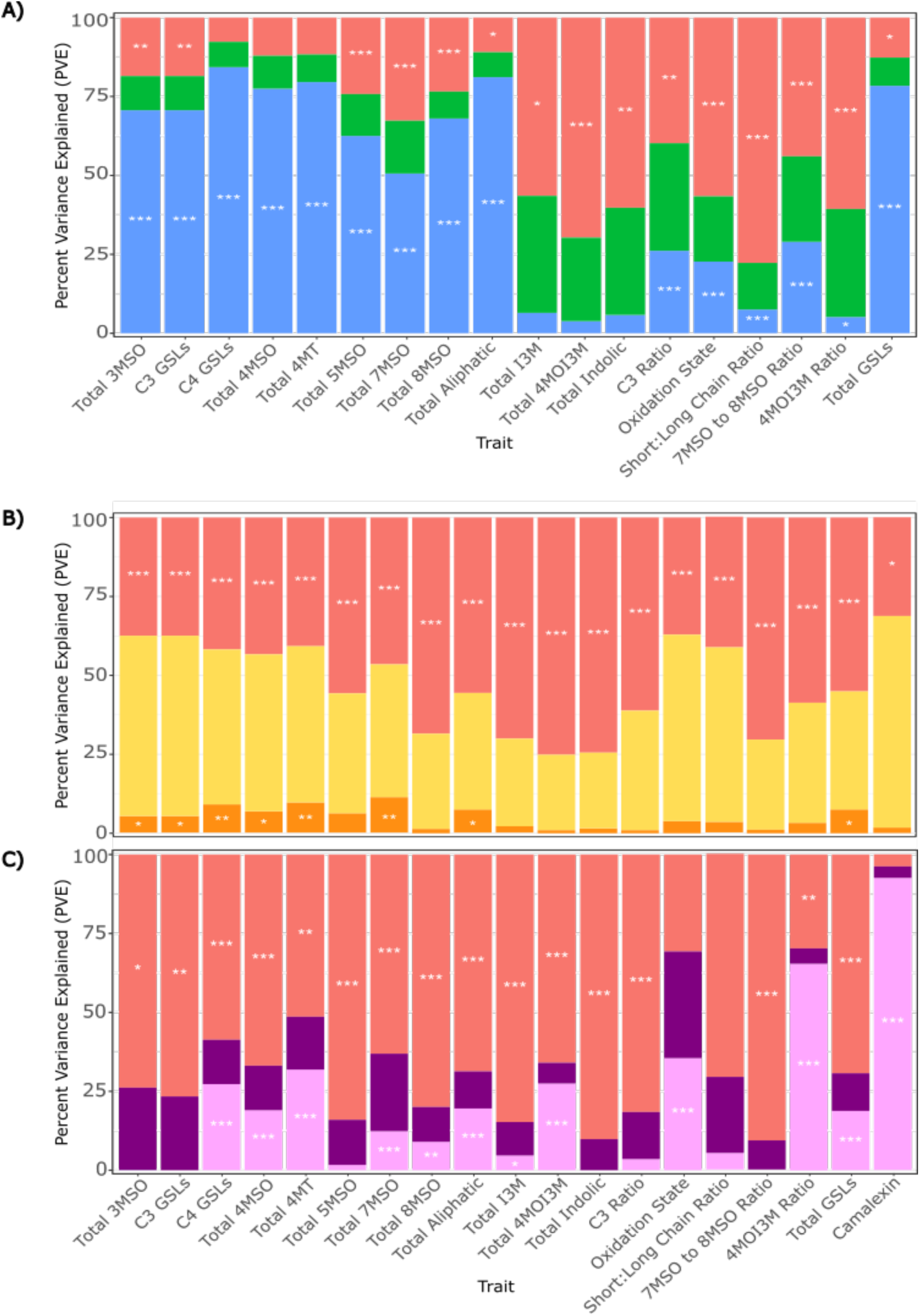
Factors modulating glucosinolates. Three LMM were used to analyze GSL data and the PVE was calculated for each factor in the models to display their relative importance. The bars display each fixed effect term from each model, and the asterisks display the significance of each term p< 0.001 "***", p< 0.01 "**", and p< 0.05 "*”. **A)** Analysis of glucosinolate across different rosette positions (inner, middle, and outer) in each genotype (genotype x rosette position). The fixed effects are represented as listed: genotype (red), genotype x rosette position (Green) and rosette position (Blue). **B)** Isolate specific variation influencing glucosinolates. The linear model tested for genotype x isolate effects accounting for differences between the six *B. cinerea* isolates regarding GSL content and camalexin 48 HPI on the 30 genotypes. The fixed effects are represented as listed: genotype (red), genotype x isolate (Yellow) and isolate (Orange). **C)** Impact of pathogen induction on glucosinolates was tested using a third model that analyzed the genotype x treatment effect to determine if there was a metabolic response to the presence of *B. cinerea* compared to uninfected leaves at 48 HPI. The fixed effects are represented as listed: genotype (red), genotype x treatment (Purple) and treatment (Pink).

**Figure 4:**
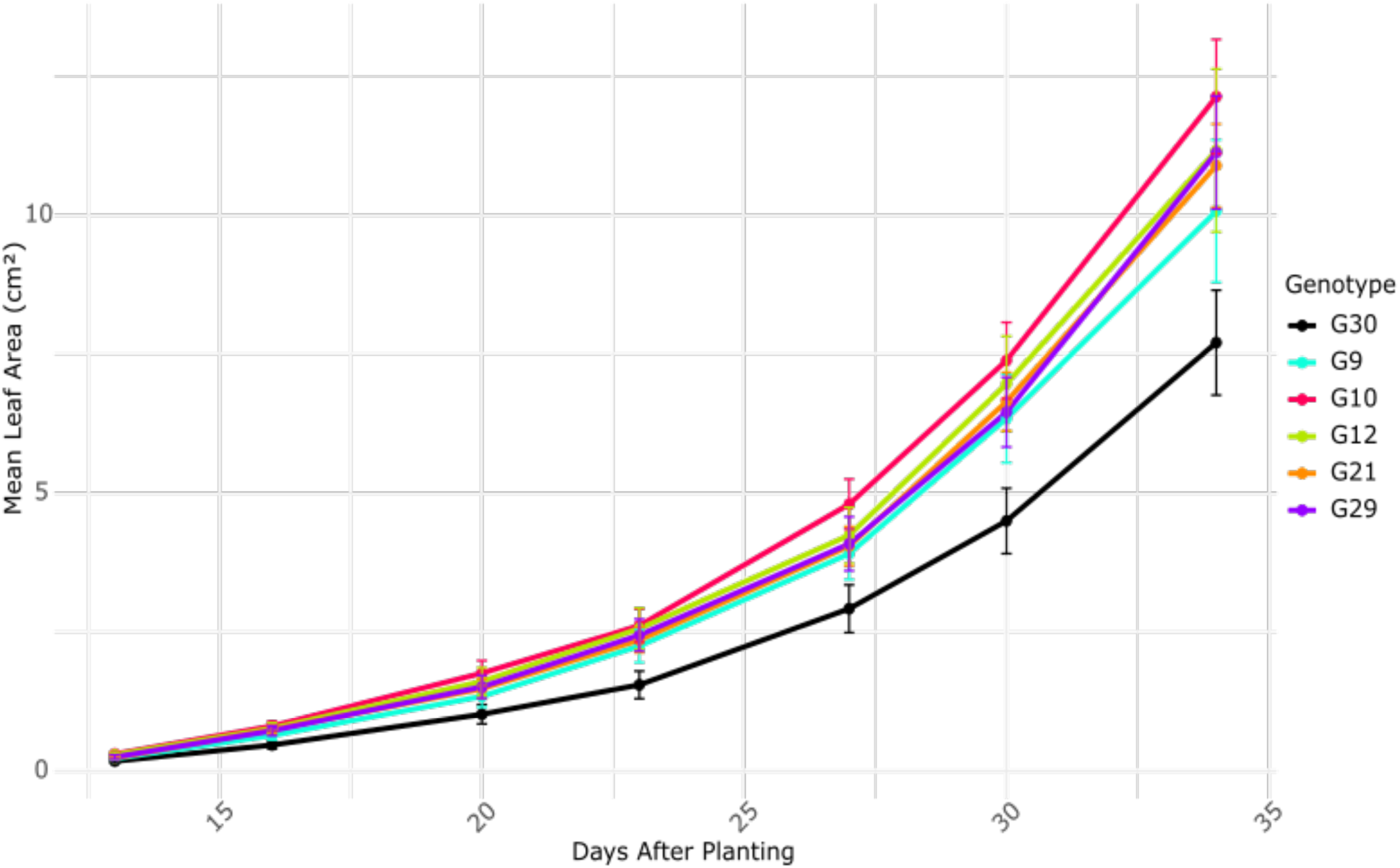
Growth curve of genotypes with disrupted cis-element motifs growing significantly faster than wild-type. Contrasts against wild-type/G30 were performed at each day to calculate the estimated leaf area (cm^2^) with FDR correction (FDR<0.05). All statistical analysis is listed in Supplemental Tables 3 and 4.

Contrasts to wild-type (G30) identified eight mutant genotypes with significant effects on at least one GSL trait. Among these, G7, with mutations in motifs 4, 5, and 7, had the strongest phenotype, with decreased levels of nearly all aliphatic GSLS across all the rosette positions, while the amount of indolic GSLs remained unaffected (Supplemental Table 3). Genotypes 10, 18, and 20 also had significant variation in aliphatic GSL accumulation but were limited to single compounds. Genotype 4 had consistently lower oxidative state and reduced short-to-long chain ratio in both the middle and outer rosettes, suggesting a potential disruption in the GSL chain elongation processes. Genotypes 23, 25, and 27 were each significant for different individual GSLs, indicating that mutations may cause subtle modulation of the pathway, rather than broad shifts in GSL biosynthesis. All the genotypes with significant GSL differences have a mutation in Motif 7 of AT3G13930 and motif 5 of AT1G54220. However, the specific effects are not fully consistent across all mutant lines. Overall GSL biosynthesis could be subtly affected due to mutated shared transcriptional control points and the individual genotypes possibly through differential penetrance or allelic variation around this common point.

### Effects of *B. cinerea* Presence and Absence on GSL and Camalexin Accumulation

To examine if cis-regulatory mutations in the mtPDC gene promoters alter pathogen-induced defense metabolism, we measured GLS and camalexin accumulation in response to *B. cinerea* infection. Defense compounds are often regulated in response to pathogens and depend on mtPDC due to its role in supplying acetyl-CoA for energy conversion and amino acid biosynthesis via TCA cycle derivatives (Kliebenstein, 2008). Thus, we hypothesized that any mutation constraining the mtPDC complex may display an enhanced phenotype after exposure to a pathogen that increases the metabolic flux towards defense metabolites. Across mutant lines, infection with *B. cinerea* significantly altered GSLs profiles, with the strongest effects observed in the indolic GSLs and camalexin, consistent with prior findings (Kliebenstein et al., 2005). Unexpectedly, the genotypes explained more variance in GSL accumulation than infection status, and no evidence for genotype x isolate interaction was detected (Figure 3B-C). This was equally true whether modeling general *B. cinerea* infection effects or isolate-specific responses (Figure 3B-C). This suggests that cis-regulatory mutations primarily alter GSL accumulation, independently of *B. cinerea* infection.

Fifteen genotypes (2, 5, 7, 8, 11, 12, 13, 16, 18, 20, 22, 23, 25, 27, and 28) showed a significant difference in at least one GSL trait in comparison to wild-type. G12 (mutated in motifs 2, 5, and 7) showed the greatest number of significant GSL contrasts to wild-type involving nearly all indolic and aliphatic GSLs. In contrast, G7 (mutated in motifs 4, 5, and 7) had an impact on only two GSLs traits in response to pathogen attack (long chain ratio and short chain ratio) even though it contained decreased levels of nearly all aliphatic GSLS across all the rosette positions with no treatment (Supplemental Table 3). Genotypes 11, 20, and 23, all mutated in motif 7, were significant for multiple indolic GSLs (Supplemental Table 3). While some genotypes shared motif mutations, the specific combinations of mutated motifs varied, resulting in diverse GSL effects. Genotypes 7 and 12 are the most relevant across all the GSL traits. These genotypes shared motif mutations 5 and 7 but also contain mutations in differing motifs. While the effects are variable, these two motifs consistently appear in genotypes with altered GSLs (Supplemental Table 3).

While genotype remained the dominant driver of GSL variation, isolate-specific effects on individual GSL traits were detected in genotypes 5, 8, 12, 13, and 24. The C3 ratio was significant in G24 only when infected with isolate 1.03.18 (Supplemental Table 3). Similarly, the production of C3 GSLS was significantly affected in G5 upon infection with Molly, C4 GSLs were affected in G12 upon infection with 1.04.04, and total 4MOI3M GSLs were affected in G8 upon infection with KernB2 (Supplemental Table 3). These significant differences from wild type in C3, C4, and 4MOI3M GSL levels suggest defensive upregulation in response to the application of a biotic stress as well as fine tune changes in metabolism dependent on genotype and condition. The significant isolate effects mostly correlated with the same genotypes and GSLs located with the *B. cinerea* absence/presence model. These results demonstrate that although genotype has the strongest influence on GSL accumulation under pathogen pressure, individual isolates were also eliciting some metabolic responses in tied to specific GSL traits, but there was no significant genotype x isolate interaction.

### Transcriptomic consequences of mtPDC cis-element variation

To test if the cis-element variation introduced into the mtPDC promoters affect the plants transcriptome, we performed RNA-sequencing on leaf tissue collected concurrently from plants grown in for the *B. cinerea* experiments. After filtering out low-quality libraries and transcripts with low read counts, a total of 13,773 transcripts were analyzed. We compared the gene expression of every mutant genotype vs. the wild-type G30 to identify differentially expressed genes in each genotype. Focusing on the specific mtPDC genes showed that overall, there was typically a subtle genotype effect across the mtPDC genes in many of the mutant genotypes. Only AT3G13930 showed a strong decrease in expression and only in some of genotypes (G2, G10 and G12) with cis-regulatory mutants within their promoters (Figure 5, Supplemental Table 3). Thus, mutations in these promoter elements do not cause strong effects on the expression of most mtPDC genes in our experimental conditions. This suggests extensive redundancy in the promoter where the other elements either compensate or dominate the expression in these conditions (Supplemental Table 3).

**Figure 5:**
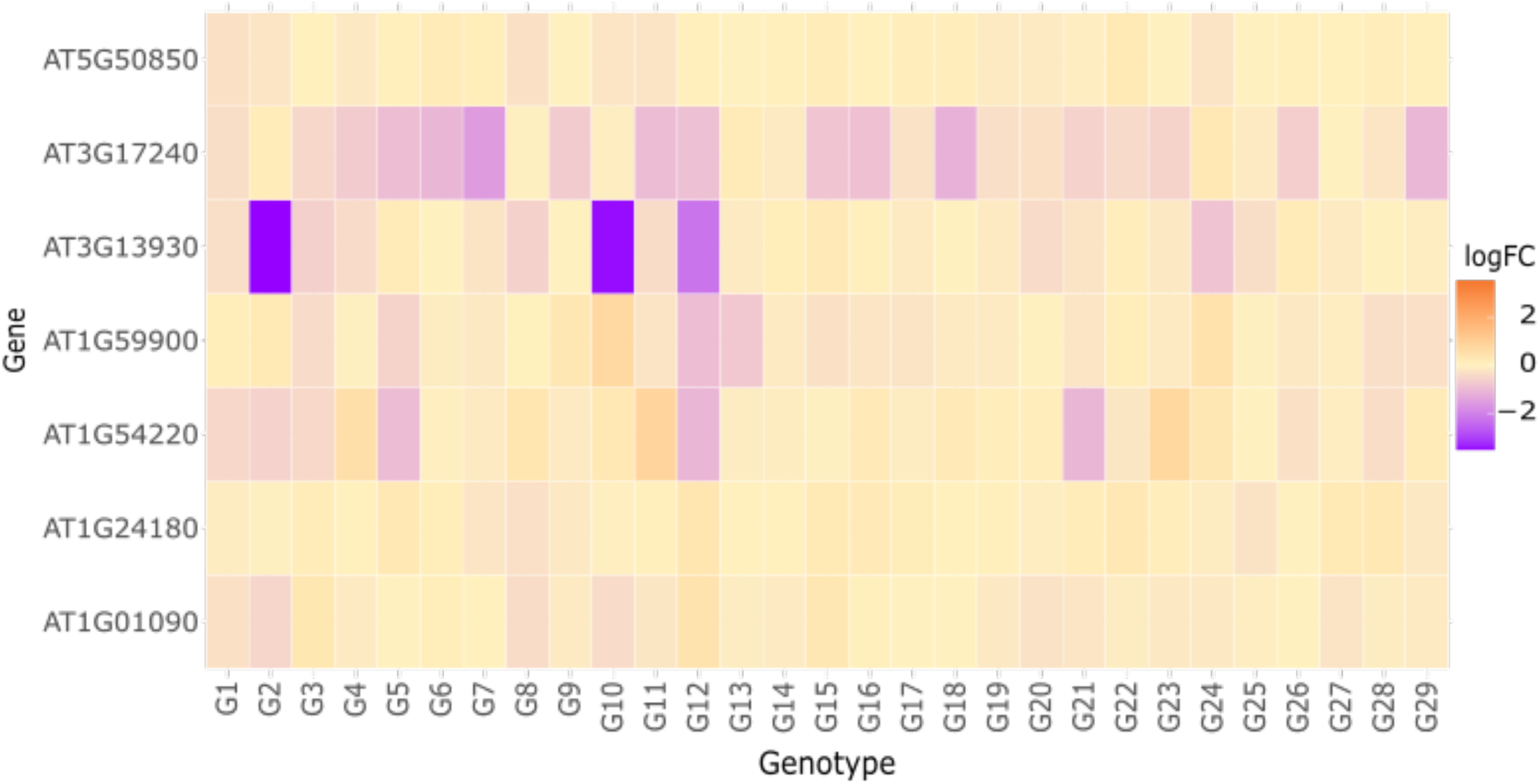
Genotype Effect on mtPDC gene expression. The Genotype model tests the mtPDC genes listed for differential expression across the genotypes in reference to wild-type/G30. Each logFC is the estimated fold change in each genotype relative to the wild type G30. These genes were filtered for the mtPDC genes targeted with CRISPR/Cas9. The color fill indicates if the genes were upregulated (positive) or downregulated (negative). The color scale was centered at zero. The significance threshold used the Benjamini-Hochberg false discovery rate (FDR) where for each genotype, a gene is significant if FDR < 0.05 vs G30.

Across all genotypes within the full transcriptome, 438 unique genes were found to be significantly differentially expressed (FDR < 0.05) in at least one genotype (G1–G29) compared to wild-type (Figure 8). To evaluate similarities in the transcripts and/or processes altered in the genotypes, we focused on GO enrichment of the genes individually altered in each genotype relative to G30. Differentially expressed pathways shared among the most genotypes included pectin, and carbohydrate catabolic processes, galacturonan metabolic processes, pollen sperm cell differentiation, microgametogenesis, and pollen development (Supplemental Figure 3, Supplemental Table 5). This suggests that many genotypes alter pathways involved in cell wall modification and reproductive development. Processes connected to cell wall modification and reproductive development are central to growth, defense and stress adaptation which could also explain why the effects of each motif mutation were spread out and variable (Chaturvedi et al., 2021; García-Angulo & Largo-Gosens, 2022; Shin et al., 2021).

Using the GO-terms, we clustered the genotypes based on shared alterations in biological processes. This identified four main clusters of genotypes with similar altered processes (Supplemental Figure 2). As expected, all genotypes are enriched in general metabolism GO terms which is also expected due to mtPDC being central to energy metabolism. Cluster 1 included genotypes with altered expression of genes involved in glycoprotein metabolic processes, microtubule processes, carbohydrate biosynthesis, glycosylation processes and amino sugar metabolic processes. Cluster 2 was enriched for GO terms related to cell growth and morphogenesis, indole glucosinolate and coumarin metabolism, protein signaling, and cellular trafficking. Four same genotypes (G12, G20, G23, and G25) that were significant for indole GSLs in the GSL *B. cinerea* experiments, were also part of this cluster (Supplemental Figure 2). Cluster 4 overlapped with clusters 1 and 2 in enriched terms but also included oxylipin biosynthetic processes. Cluster 3 is made up solely by the G2 genotype and has altered expression in genes linked to acetyl-CoA and protein and nucleotide metabolism. Interestingly, acetyl-coA is the precursor for the mtPDC complex and its GO terms only link to the G2 mutant which also contains the most promoter edits (motif 2, 3, 5, 7 and 9) within the mtPDC subunit genes (Figure 1). This result suggests that those multiple cis-element mutations are beginning to impact mtPDC pathway function and further studies may require broader mutagenesis of promoter regions to further perturb this pathway (Figure 1 and 2, Supplemental Figure 2, Supplemental Table 5). Importantly, the number of mutated motifs alone does not fully explain the clustering. G3 also harbors five promoter mutations, yet it is not grouped with G2, likely due to differences in the specific motifs or genes targeted (Supplemental Figure 2). This suggests that both the identity and context of cis-regulatory mutations contribute to transcriptomic outcomes.

### RNA Sequencing Motif Model

As common transcriptomic effects clustered the cis-regulatory mutant genotypes, we tested if these effects are created by specific promoter motifs. To do this, we compared each transcript’s expression between mutant and wild-type samples by nesting the genotypes into either wild-type or mutant for each motif and explicitly modeling the motif term across all genotypes (Table 1, Figure 6, and Supplemental Table 1). Motifs 8 and 9 were dropped from this model because they don’t have enough levels to generate the contrasts and comparisons. Investigating the mtPDC genes showed that the presence of mutations in motif 3 within AT1G48030 and motif 5 in AT1G54220 led to a significant decrease in AT3G13930 (Supplemental Table 3). Interestingly, both motif mutations had their strongest transcriptional effect on a different gene than the gene’s promoter in which they located. This suggests the existence of compensatory regulation mechanisms occurring within the mtPDC complex set of genes (Figure 6).

**Figure 6:**
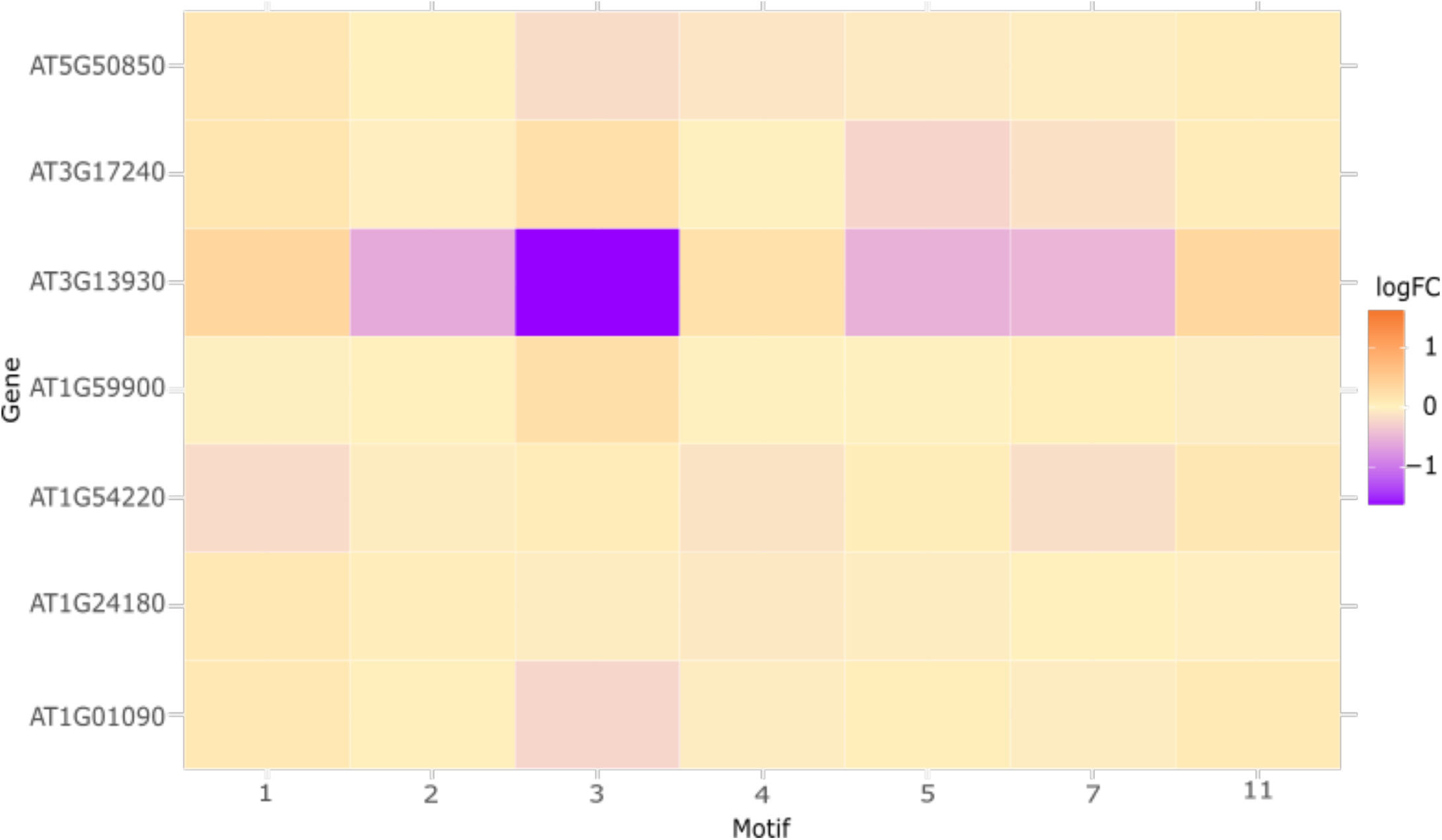
Promoter Motif Effect on mtPDC Gene Expression (Log FC). For each mtPDC transcript, the different genotypes with mutations in specific motifs in specific promoters were combined into a linear model to test if the promoter motif mutations in specific cis-regulatory elements (e.g., ATHB sites) altered transcripts in common. This model tests the motif effect across genotypes, while adjusting for genotype-specific backgrounds (i.e. the genotype term is nested within motif mutation wild-type vs. mutant Expression ∼ Motif/Genotype). This adjusts for the genotype specific background and selects for motif associated DEGs in mtPDC specific genes. Motifs 8 and 9 were dropped due to having only one mutant genotype level. The significance threshold used the Benjamini-Hochberg false discovery rate (FDR) where for each motif a gene in post-hoc tests against wild-type at < 0.05.

We next expanded this approach to assay the entire transcriptomic dataset to test if the cis-regulatory motifs have overlapping or distinct expression effects on the transcriptome. This analysis showed that each motif had a unique transcriptomic signature, with few overlaps (Figure 7, Supplemental Figure 3, Supplemental Table 6). Motif 2 (pAT1G1090, subunit E1), motif 5 (pAT1G54220, subunit E2), and motif 7 (pAT3G13930, subunit E2) are all enriched for changes in cellular oxygen response and hypoxia. The upset plots of the top genes located per motif also confirmed these overlaps between these three motifs (Figure 7, Supplemental Table 6). In contrast, some motifs are associated with different processes and have little overlap. Mutations in Motif 3 (pAT1G48030, subunit E3) are associated with alterations in photosynthetic response pathways, whereas Motif 1 (pAT1G01090, subunit E1) is enriched for genes related to water, fatty acid, fungi, and jasmonic acid (Supplemental Table 6). Regarding metabolism, Motif 3 was associated with GO terms connected to sulfur compound metabolic processes, amide metabolic processes, and acetyl-CoA metabolic processes in association with mtE2-2, whereas Motif 5 was associated with monocarboxylic acid metabolic processes, connected to mtE2-2 (the mtPDC E2 subunit) (p<0.05) (Supplemental Table 6). Motif 11 (pAT3G52200, subunit E2) also stood apart, associating with metal and iron homeostasis pathways. Regarding cellular localization, Motif 3 was almost exclusively associated with photosynthetic and chloroplast-related GO terms with minimal overlap. Many of the significantly affected pathways are connected to mtPDC function. Each mtPDC gene was screened via the BAR eFP Browser on TAIR to search for candidate stress experiments, and almost every mtPDC gene was sensitive to selenium and sulfur. As a monocarboxylic acid, pyruvate is the first compound from glycolysis that the mtPDC processes to create acetyl-CoA. These observations suggest that individual cis-regulatory motifs hold partial co-regulation attributable to being bound by the same TF. However, variation in biological context or environmental cues may drive unique combinatorial interactions, allowing for context-dependent regulation of specific mtPDC subunits.

**Figure 7:**
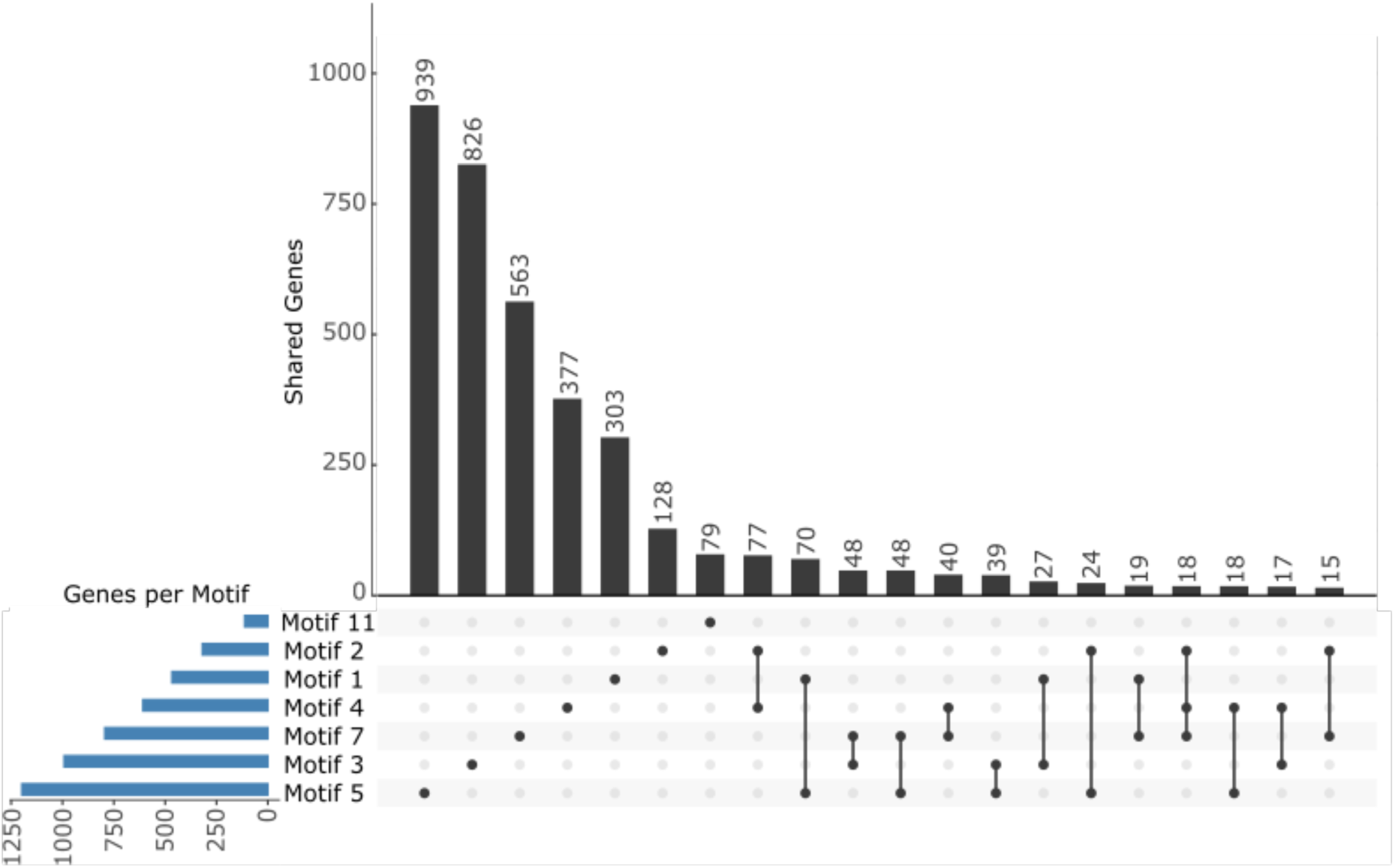
Effects on gene expression of motif disruptions in seven cis-element motifs in the mtPDC. An upset plot shows the number of transcripts in the transcriptome significantly altered by the promoter motif mutations in comparison to the wild-type motif. The bars to the left show the total number of genes unique to each motif and the intersection bars display how many genes are shared between different motif combinations.

## Discussion

Previous analysis showed that ATHB34 TF mutants had significant effects on a Glucosinolates, Growth, hypocotyl length, and root length (Tang et al., 2021). This is striking considering that ATHB34 is one of 48 homeodomain-leucine zipper (HD-Zip) genes in *A. thaliana* (Roodbarkelari & Groot, 2017). In contrast to the effects of a mutation of a single ATHB34 TF, our cis-element mutations had more subtle effects. The mutants with the most mutations (G2 and G3 both with a different combination of 5 motif mutations total) did not have dramatic effects on any of the traits measured (Figure 8). The motifs had more of a combination of specific and overlapping effects suggesting the beginning of modular transcriptomic consequences surrounding the mtPDC complex. When measuring a common trait, GSL, the effects of the cis-element motifs changed across developmental and environmental perturbations (Supplemental Table 3). A bigger array of traits and environments would be required to measure the full effects of these motif mutations. The subtle and conditional effects combine to indicate that there may be sufficient redundancy or compensatory mechanisms to buffer against changes in individual cis-elements within individual promoters up to combinatorial effects across cis-elements spread multiple genes for the enzyme complex. Supporting this was the finding that the target genes were only subtly affected by mutating the ATHB motif and one specific gene in the complex was altered in response to mutations in several motifs (Figures 5 and 6).

**Figure 8:**
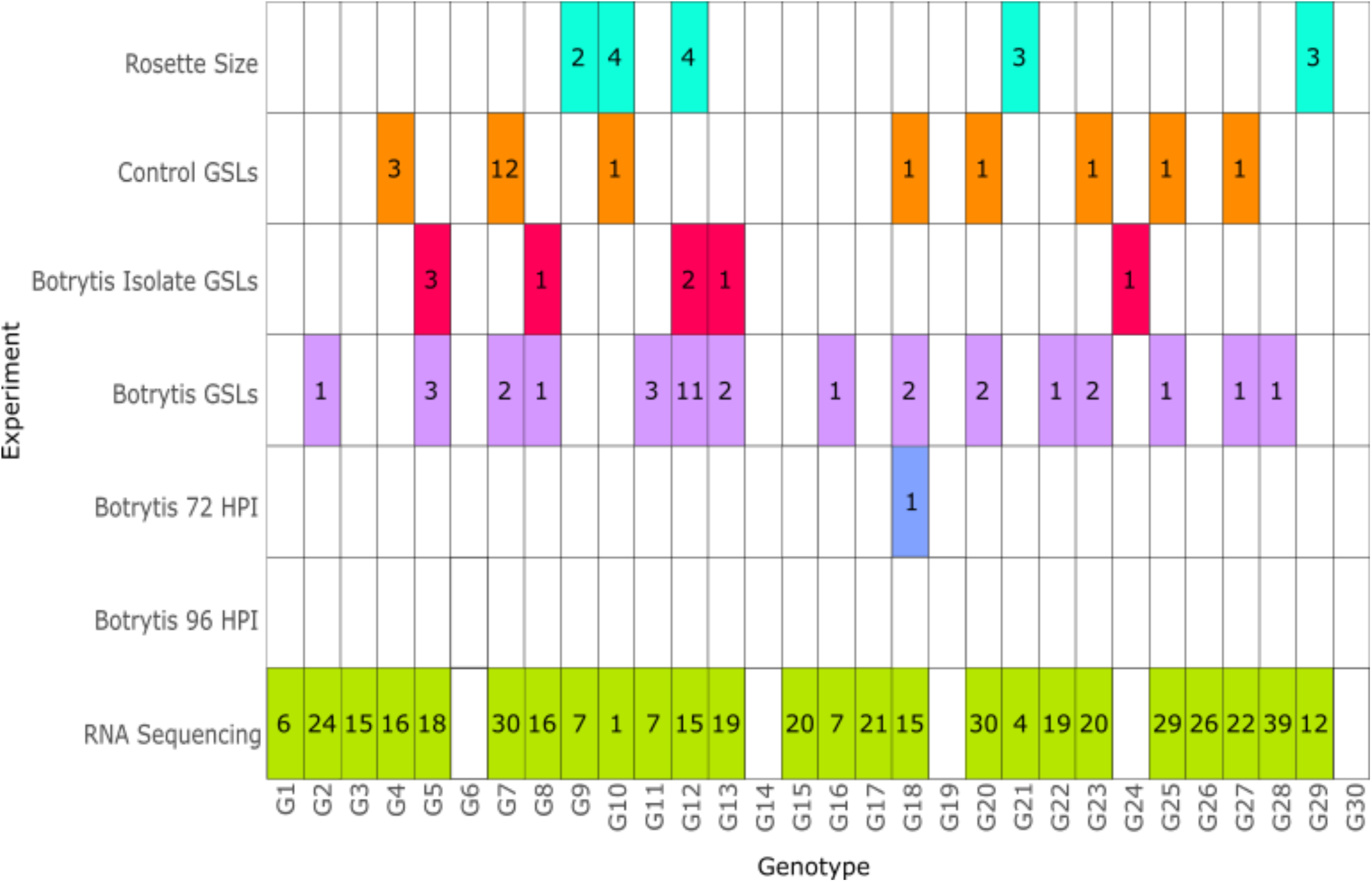
Number of Significant Genotypes Effects Linked to Each Genotype in Each Experiment The number of significant genotype effects across all the models and experiments was summarized for each genotype. Different colors represent the different experiments tested and the numbers in each box represent the number of times a genotype was significant in each experimental model, for every trait tested within each model. The RNA sequencing genotypes were filtered for all the significant GO terms located under the Genotype model.

Complicating this analysis was an inability to identify mutations in several of the motifs even when screening a large collection of progeny. Thus, it is possible that the motifs we identified mutations in are the ones with buffering properties while the motifs where we could not find mutations do not have similar levels of buffering and mutations within them are more deleterious and/or lethal. Interestingly, this differential buffering seemed to be linked to a specific promoter than to the number of ATHB motifs per promoter. This possibility would suggest that the promoters/genes even within an enzyme complex are not equivalent and that there is modularity differentiation across the genes encoding subunits for a single enzyme complex. Supporting this idea is that the transcriptomic effects of each motif showed a blend of unique effects and overlap albeit with no consistent effect of all motif mutations (Figure 7). This extended to the growth and defense measurements where the common motifs differed across traits being measured. This modularity, even within the genes encoding an enzyme complex, could indicate that the complex’s function may be finely tuned. Developing a system to force mutations in the motifs where we could not identify mutations would be needed to ascertain if these have stronger phenotypic consequences to help ascertain the modular structure and if there is differential buffering across promoters.

Overall, this analysis showed that the cis-element motif mutations within the genes for a single enzyme complex displayed distinct phenotypes illuminating the potential for buffering and modularity that appears to connect to each individual promoter. Future work is needed to test how these observations may link to metabolic pathways that do not function as enzyme complexes to test if the complex structure itself is providing some of the buffering. Blending cis-element motif mutations with associated TF mutations is also necessary to test how a TF gene family interacts with the cis-elements within a metabolic module to regulate plant growth and defense.

## Acknowledgments

We would like to thank our lab members Ritu Singh and Anna Jo Muhich, our lab technician Cloe Tom, and our undergrads Alina Stratulea, Tiffany Ng, Jaisy Huang, Richa Kakde and Benedict-Gillia Palma for help with the set up and screening efforts during these experiments.

---

**Supplemental Figure 1:**
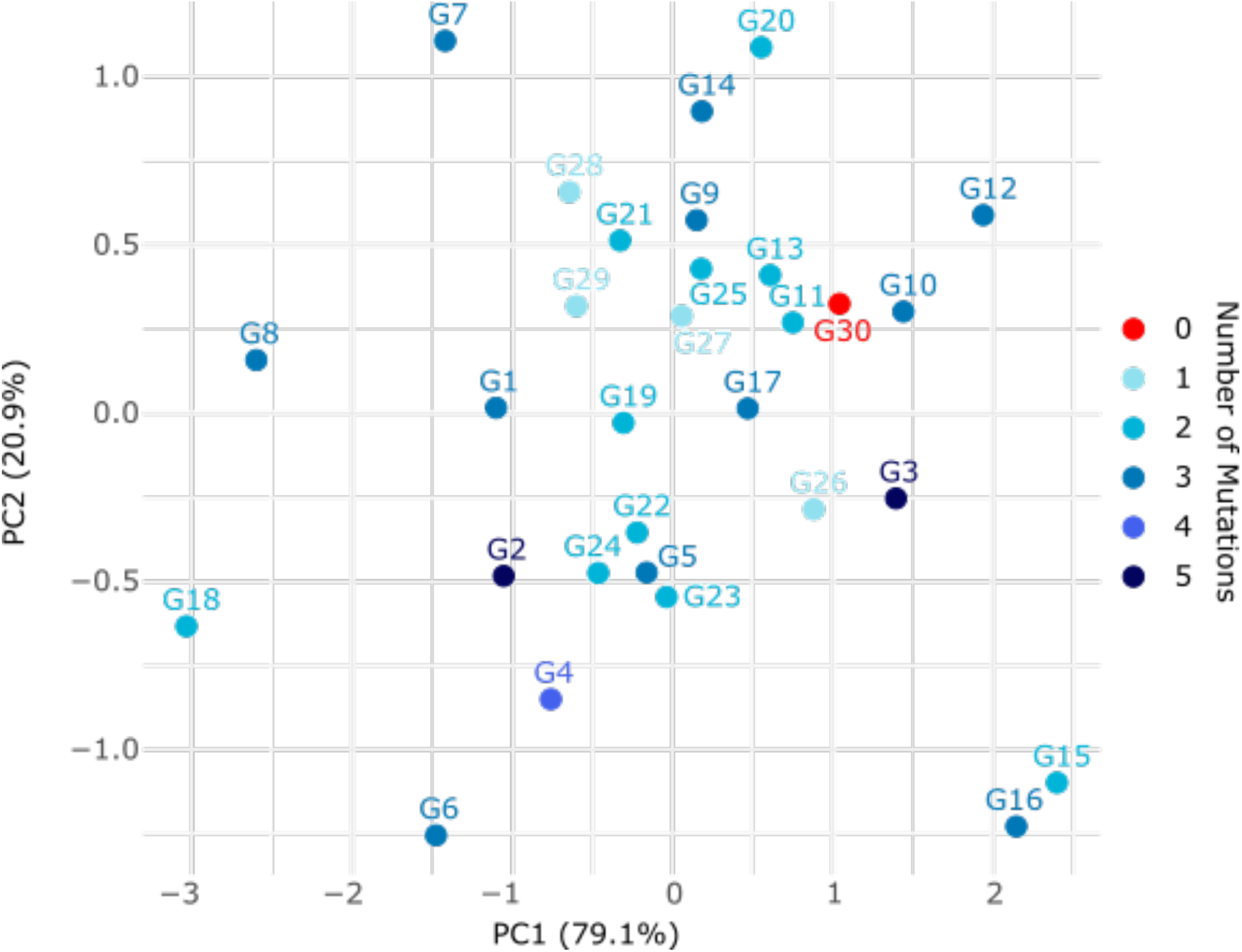
PCA of Genotype EMMeans across both 72 and 96 HPI *B. cinerea* samples. PCA was performed on mutation profiles on the genotype EMMeans for each *B. cinerea* isolate lesion formation at both the 72 and 96 HPI timepoints. The number of promoter motif mutations in each genotype is colored, and each genotype is labeled from G1 through G30/wild-type.

**Supplemental Figure 2:**
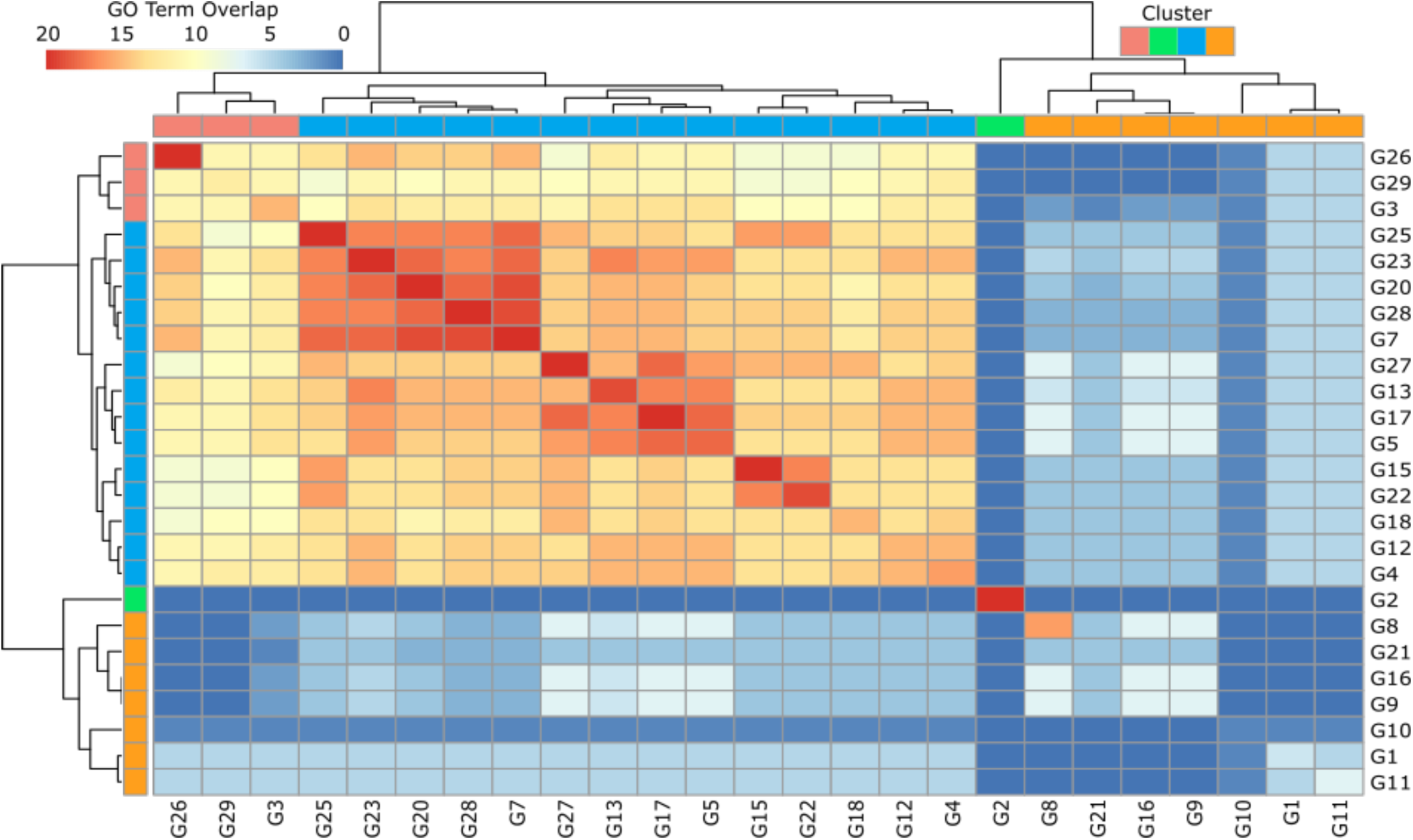
Top 20 (p<0.05) GO terms shared between promoter motif mutant genotypes vs. G30/wild-type utilizing 13,773 transcripts analyzed with the genotype model. The Euclidean distance between rows was calculated for hierarchical clustering. Red squares display many shared GO terms between genotypes and blue squares display few shared GO terms. Each genotype is mirrored from the middle for comparison.

**Supplemental Figure 3:**
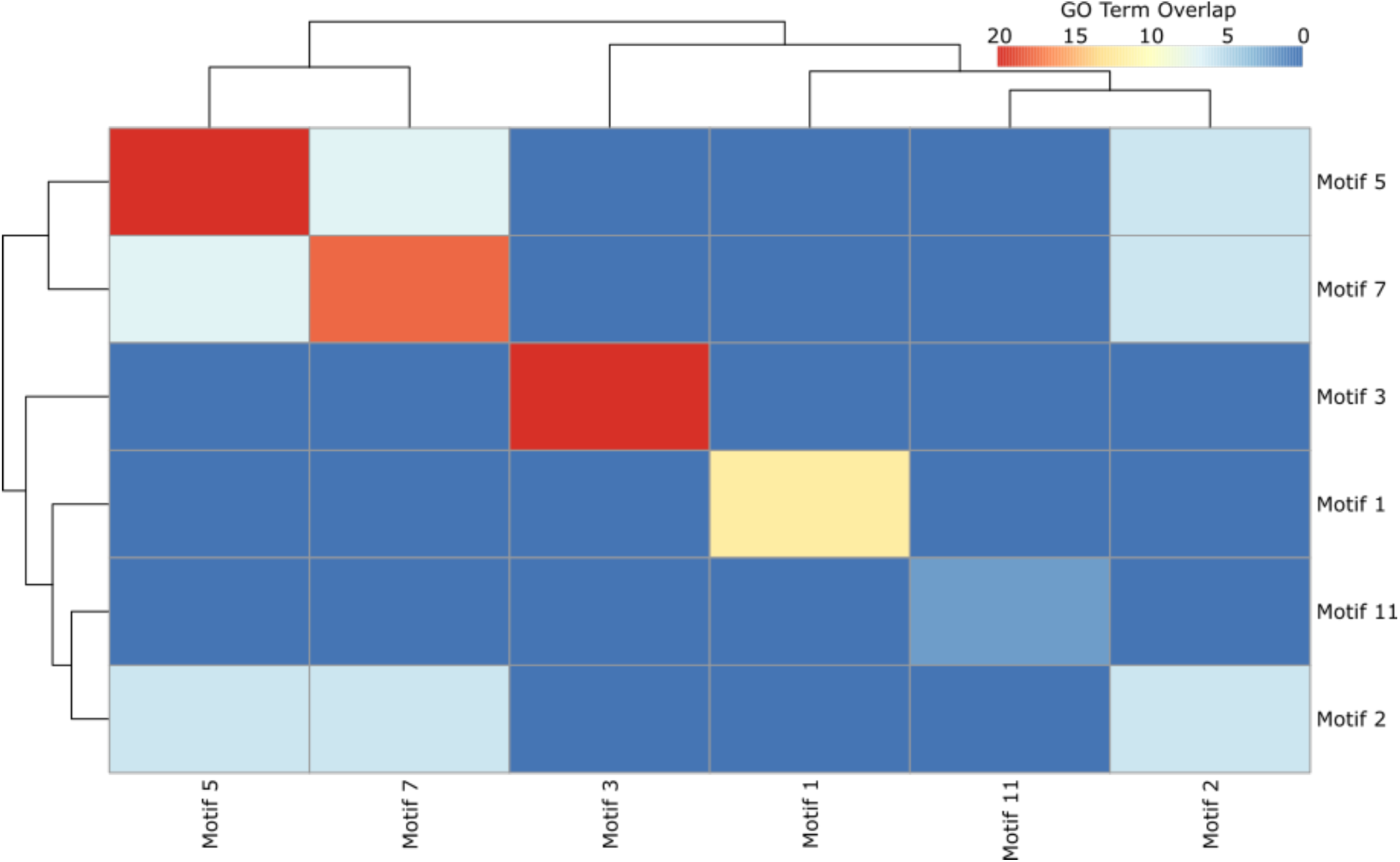
Top 20 (p<0.05) GO terms shared between shared Motif mutants vs. wild-type utilizing 13,773 transcripts analyzed with the genotype model. The Euclidean distance between rows was calculated for hierarchical clustering. Red squares display many shared GO terms between Motifs and blue squares display few shared GO terms. Each motif is mirrored from the middle for comparison.

Supplemental Table 1: Mutation Matrix; The first column lists the 29 mutant genotypes and G30/wild type. The columns represent the 13 ATHB34 motifs 1 through 13 across the different mtPDC promoters (i.e. pAT…). The specific type of mutation identified in each motif in each genotype is described. 1,3,5,30 bp insertion/deletions have successfully disrupted the ATTA motif. If a mutation is listed as "close" the gRNA was able to edit near the ATTA motif site and cause a frameshift mutation to alter the codon next to the target site.

Supplemental Table 2: Compiled ANOVA; Compiled ANOVA tables from all the linear mixed models described in the methods are provided. “Experiment” refers to the independent variable being tested (rosette growth, 72HPI and 96HPI DLA genotype model, the DLA interaction model between genotype and isolate, the GSL models (inner/middle/outer rosettes, and the *B. cinerea* isolate specific mode, and the presence/absence model). “Rosette Growth Days After Planting” is from the model leaf_Area_cm² ∼ genotype x days after planting. The experiment was run three times and the stats from all, E1, E2, and E3 are listed. There was only one main effect (genotype). The specific date from day after planting (DAP) is listed under “date”. 72 and 96 DLA Geno Model refers to the genotype model from the Detached Leaf Assays (DLA) (genotype as the fixed effect). The 72 and 96 DLA genotype x isolate Model refers to the DLA interaction model with genotype x isolate. All the HPLC models refer to the GSL analysis (genotype x rosette position, genotype x isolate and genotype x treatment); fixed effects tested are under the “effect” column and specific GSL trait is under the “Trait” column. The “Sum Sq” = sum of squares, “Mean Sq” = Mean Square, “NumDF” = numerator degrees of freedom, “DenDF” = denominator degrees of freedom, “F value” = F-statistic, “Pr(>F)” = p-value.

Supplemental Table 3: Complied Contrasts; All pairwise contrasts of each mutant to the G30/wild-type are provided for each of the linear models described. “Experiment” refers to the independent variable being tested (rosette growth, 72HPI and 96HPI DLA genotype model, the DLA interaction model between genotype and isolate, the GSL models (inner/middle/outer rosettes, and the *B. cinerea* isolate specific mode, and the presence/absence model). “Contrast” = comparison between levels. “Estimate” = Estimated difference. “SE” = standard error. “DF” = degrees of freedom. “t.ratio” = t-statistic. “p.value” = p-value. The p-values are presented as the FDR adjusted values within each model. “Rosette Growth Days After Planting” is from the model Leaf_Area_cm² ∼ genotype x days after planting. The experiment was run three times and the stats from the compiled experiments are listed under “trait” for each genotype (listed under contrast) and the DAP (under “trait”). 72 and 96 DLA Geno Model refers to the genotype model from the DLA (genotype as the fixed effect). There are no stats from the interaction model because the ANOVA showed no significant interaction between genotype and isolate. The HPLC Rosette GSL model tested genotype x rosette position, and the genotype contrast are under “contrast”, specific GSL trait is under “trait” and the inner, outer, and middle refers to “rosette position”. The HPLC Isolate model is similar except instead of testing rosette position it tested genotype x isolate and each of the *B. cinerea* isolates are listed under “Isolate”. The HPLC *B. cinerea* Y/N model tested genotype x treatment (i.e. yes or no *B. cinerea*). Yes or no are under the “Isolate” column to denote presence and absence of any isolates.

Supplemental Table 4: EMMeans; All EMMeans for each trait for each mutant and wild-type are provided. “EMMean” = estimated marginal mean. “SE” = Standard Error. “DF” = degrees of freedom. “Lower.CL” = Lower confidence Limit. “Upper.CL” = Upper confidence limit. Experiment lists the model, Genotype is G1-G30, DAP is for the rosette DAP model, Isolate / rosette position / metabolite traits are all for the HPLC models. Isolate also is listed for the DLA models. Lesion size is measured in mm^2^, GSLs are in umol, and leaf area across DAP is in cm^2^. “Rosette Growth Days After Planting” is from the model Leaf_Area_cm² ∼ genotype x days after planting. The experiment was run three times and the stats from the compiled experiments are listed under “trait” for each genotype (listed under contrast) and the DAP (under “trait”). 72 and 96 DLA Geno Model refers to the genotype model from the DLA (genotype as the fixed effect). There are no stats from the interaction model because the ANOVA showed no significant interaction between genotype and isolate. The HPLC Rosette GSL model tested genotype x rosette position, and the genotype contrast are under “contrast”, specific GSL trait is under “trait” and the inner, outer, and middle refers to “rosette position”. The HPLC Isolate model is similar except instead of testing rosette position it tested genotype x isolate and each of the *B. cinerea* isolates are listed under “Isolate”. The HPLC *B. cinerea* Y/N model tested genotype x treatment (i.e. yes or no *B. cinerea*). Yes or no are under the “Isolate” column to denote presence and absence of any isolates.

Supplemental Table 5: All GO terms and all contrasts to G30 (Genotype Model). The pairwise contrasts generated via Limma were used for GO enrichment. This was performed by filtering genes with adjusted p-values < 0.05, mapped *A. thaliana* gene identifiers (TAIR) to Entrez IDs and conducted biological process (BP) enrichment using the clusterProfiler package in R. The output from GO enrichment analysis as follows: ID: GO Term ID from gene ontology database, Description: GO term metabolic/biological process or cellular component summary, GeneRatio: ratio of significant input genes annotated with the GO term, BgRatio: Ratio of background genes, Rich Factor: calculated as GeneRatio/BgRatio, Count: number of genes in the input associated with the GO term, Contrast: experimental contrast being tested, Signficant: tests if GO term passes FDR < 0.05.

Supplemental Table 6: All GO terms and all contrasts to G30 (Motif Model). The pairwise contrasts generated via Limma were used for GO enrichment. This was performed by filtering genes with adjusted p-values < 0.05, mapped *A. thaliana* gene identifiers (TAIR) to Entrez IDs and conducted biological process (BP) enrichment using the clusterProfiler package in R. The output from GO enrichment analysis as follows: ID: GO Term ID from gene ontology database, Description: GO term metabolic/biological process or cellular component summary, GeneRatio: ratio of significant input genes annotated with the GO term, BgRatio: Ratio of background genes, Rich Factor: calculated as GeneRatio/BgRatio, Count: number of genes in the input associated with the GO term, Contrast: experimental contrast being tested, Signficant: tests if GO term passes FDR < 0.05.

## References

Altschul, S. F., Gish, W., Miller, W., Myers, E. W., & Lipman, D. J. (1990). Basic local alignment search tool. Journal of Molecular Biology, 215(3), 403–410. 10.1016/S0022-2836(05)80360-2

Brown, P. D., Tokuhisa, J. G., Reichelt, M., & Gershenzon, J. (2003). Variation of glucosinolate accumulation among different organs and developmental stages of Arabidopsis thaliana. Phytochemistry, 62(3), 471–481. 10.1016/S0031-9422(02)00549-6

BTI Curriculum Development Projects in Plant Biology. (2015). ImageJ Measurement Protocol.

Butow, R. A. (1999). A Transcriptional Switch in the Expression of Yeast Tricarboxylic Acid Cycle Genes in Response to a Reduction or Loss of Respiratory Function †. 19(10), 6720–6728.

Campbell, N. R., Harmon, S. A., & Narum, S. R. (2015). Genotyping-in-Thousands by sequencing (GT-seq): A cost effective SNP genotyping method based on custom amplicon sequencing. Molecular Ecology Resources, 15(4), 855–867. 10.1111/1755-0998.12357

Caseys, C., Muhich, A. J., Vega, J., Ahmed, M., Hopper, A., Kelly, D., Kim, S., Madrone, M., Plaziak, T., Wang, M., & Kliebenstein, D. J. (2024). Leaf abaxial and adaxial surfaces differentially affect plant-fungal pathogen interactions (Vol. 27, Issue 2, pp. 635–637). 10.1101/2024.02.13.579726

Caseys, C., Shi, G., Soltis, N., Gwinner, R., Corwin, J., Atwell, S., & Kliebenstein, D. J. (2021). Quantitative interactions: The disease outcome of Botrytis cinerea across the plant kingdom. G3: Genes, Genomes, Genetics, 11(8). 10.1093/g3journal/jkab175

Čermák, T., Curtin, S. J., Gil-Humanes, J., Čegan, R., Kono, T. J. Y., Konečná, E., Belanto, J. J., Starker, C. G., Mathre, J. W., Greenstein, R. L., & Voytas, D. F. (2017). A multipurpose toolkit to enable advanced genome engineering in plants. Plant Cell, 29(6), 1196–1217. 10.1105/tpc.16.00922

Chaturvedi, P., Wiese, A. J., Ghatak, A., Záveská Drábková, L., Weckwerth, W., & Honys, D. (2021). Heat stress response mechanisms in pollen development. New Phytologist, 231(2), 571–585. 10.1111/nph.17380

Doench, J. G., Fusi, N., Sullender, M., Hegde, M., Vaimberg, E. W., Donovan, K. F., Smith, I., Tothova, Z., Wilen, C., Orchard, R., Virgin, H. W., Listgarten, J., & Root, D. E. (2016). Optimized sgRNA design to maximize activity and minimize off-target effects of CRISPR-Cas9. Nature Biotechnology, 34(2), 184–191. 10.1038/nbt.3437

Duncan, O., Taylor, N. L., Carrie, C., Eubel, H., Kubiszewski-Jakubiak, S., Zhang, B., Narsai, R., Harvey Millar, A., & Whelan, J. (2011). Multiple lines of evidence localize signaling, morphology, and lipid biosynthesis machinery to the mitochondrial outer membrane of Arabidopsis. Plant Physiology, 157(3), 1093–1113. 10.1104/pp.111.183160

Ferrari, S., Galletti, R., Denoux, C., De Lorenzo, G., Ausubel, F. M., & Dewdney, J. (2007). Resistance to Botrytis cinerea induced in arabidopsis by elicitors is independent of salicylic acid, ethylene, or jasmonate signaling but requires PHYTOALEXIN DEFICIENT3. Plant Physiology, 144(1), 367–379. 10.1104/pp.107.095596

García-Angulo, P., & Largo-Gosens, A. (2022). Plant Cell Wall Plasticity under Stress Situations. Plants, 11(20), 10–11. 10.3390/plants11202752

Gaudinier, A., Tang, M., & Kliebenstein, D. J. (2015). Transcriptional networks governing plant metabolism. Current Plant Biology, 3–4, 56–64. 10.1016/j.cpb.2015.07.002

Hoang, X. L. T., Nhi, D. N. H., Thu, N. B. A., Thao, N. P., & Tran, L.-S. P. (2017). Transcription Factors and Their Roles in Signal Transduction in Plants under Abiotic Stresses. Current Genomics, 18(6), 483–497. 10.2174/1389202918666170227150057

Hunziker, P., Lambertz, S. K., Weber, K., Crocoll, C., Halkier, B. A., & Schulz, A. (2021). Herbivore feeding preference corroborates optimal defense theory for specialized metabolites within plants. Proceedings of the National Academy of Sciences of the United States of America, 118(47), 1–6. 10.1073/pnas.2111977118

Ihmels, J., Bergmann, S., & Barkai, N. (2004). Defining transcription modules using large-scale gene expression data. Bioinformatics, 20(13), 1993–2003. 10.1093/bioinformatics/bth166

Ihmels, J., Friedlander, G., Bergmann, S., Sarig, O., Ziv, Y., & Barkai, N. (2002). Revealing modular organization in the yeast transcriptional network. Nature Genetics, 31(4), 370–377. 10.1038/ng941

Invitrogen. (2006). One Shot ® Mach1 ^TM^ -T1. Cat. No. C8620-03, October, 1–2.

Jensen, L. M., Jepsen, H. S. K., Halkier, B. A., Kliebenstein, D. J., & Burow, M. (2015). Natural variation in cross-talk between glucosinolates and onset of flowering in Arabidopsis. Frontiers in Plant Science, 6(September), 1–10. 10.3389/fpls.2015.00697

Katz, E., Li, J.-J., Jaegle, B., Ashkenazy, H., Abrahams, S. R., Bagaza, C., Holden, S., Pires, C. J., Angelovici, R., & Kliebenstein, D. J. (2021). Genetic variation, environment and demography intersect to shape Arabidopsis defense metabolite variation across Europe. ELife, 10, 1–25. 10.7554/elife.67784

Kliebenstein, D. J. (2008). A role for gene duplication and natural variation of gene expression in the evolution of metabolism. PLoS ONE, 3(3). 10.1371/journal.pone.0001838

Kliebenstein, D. J., Gershenzon, J., & Mitchell-Olds, T. (2001). Comparative quantitative trait loci mapping of aliphatic, indolic and benzylic glucosinolate production in Arabidopsis thaliana leaves and seeds. Genetics, 159(1), 359–370. 10.1093/genetics/159.1.359

Kliebenstein, D. J., Kroymann, J., Brown, P., Figuth, A., Pedersen, D., Gershenzon, J., & Mitchell-Olds, T. (2001). Genetic control of natural variation in arabidopsis glucosinolate accumulation. Plant Physiology, 126(2), 811–825. 10.1104/pp.126.2.811

Kliebenstein, D. J., Lambrix, V. M., Reichelt, M., Gershenzon, J., & Mitchell-Olds, T. (2001). Gene duplication in the diversification of secondary metabolism: Tandem 2-oxoglutarate-dependent dioxygenases control glucosinolate biosynthesis in arabidopsis. Plant Cell, 13(3), 681–693. 10.1105/tpc.13.3.681

Kliebenstein, D. J., Rowe, H. C., & Denby, K. J. (2005). Secondary metabolites influence Arabidopsis/Botrytis interactions: Variation in host production and pathogen sensitivity. Plant Journal, 44(1), 25–36. 10.1111/j.1365-313X.2005.02508.x

Kumar, R., Ichihashi, Y., Kimura, S., Chitwood, D. H., Headland, L. R., Peng, J., Maloof, J. N., & Sinha, N. R. (2012). A high-throughput method for Illumina RNA-Seq library preparation. Frontiers in Plant Science, 3(AUG), 1–10. 10.3389/fpls.2012.00202

Lan, X., Farnham, P. J., & Jin, V. X. (2012). Uncovering transcription factor modules using one- and three-dimensional analyses. Journal of Biological Chemistry, 287(37), 30914–30921. 10.1074/jbc.R111.309229

Li, B., Gaudinier, A., Tang, M., Taylor-Teeples, M., Nham, N. T., Ghaffari, C., Benson, D. S., Steinmann, M., Gray, J. A., Brady, S. M., & Kliebenstein, D. J. (2014). Promoter-based integration in plant defense regulation. Plant Physiology, 166(4), 1803–1820. 10.1104/pp.114.248716

Li, X., Yu, J., Zhou, W., Yan, F., Lin, C., & Tao, Z. (2025). A homeobox transcription factor HB34 suppresses jasmonic acid biosynthesis but promotes the expression of growth-related genes to balance plant immunity and growth in Arabidopsis. Plant Communications, 6(9), 101429. 10.1016/j.xplc.2025.101429

Liu, H., Ding, Y., Zhou, Y., Jin, W., Xie, K., & Chen, L. L. (2017). CRISPR-P 2.0: An Improved CRISPR-Cas9 Tool for Genome Editing in Plants. Molecular Plant, 10(3), 530–532. 10.1016/j.molp.2017.01.003

Logemann, E., Birkenbihl, R. P., Ülker, B., & Somssich, I. E. (2006). An improved method for preparing Agrobacterium cells that simplifies the Arabidopsis transformation protocol. Plant Methods, 2(1), 1–5. 10.1186/1746-4811-2-16

Melo, D., Porto, A., Cheverud, J. M., Marroig, G., & Louis, S. (2017). Modularity : genes, development and evolution. 463–486. 10.1146/annurev-ecolsys-121415-032409.Modularity

Meyer, R. C., Steinfath, M., Lisec, J., Becher, M., Witucka-Wall, H., Törjék, O., Fiehn, O., Eckardt, Ä., Willmitzer, L., Selbig, J., & Altmann, T. (2007). The metabolic signature related to high plant growth rate in Arabidopsis thaliana. Proceedings of the National Academy of Sciences of the United States of America, 104(11), 4759–4764. 10.1073/pnas.0609709104

Obayashi, T., Kinoshita, K., Nakai, K., Shibaoka, M., Hayashi, S., Saeki, M., Shibata, D., Saito, K., & Ohta, H. (2007). ATTED-II: A database of co-expressed genes and cis elements for identifying co-regulated gene groups in Arabidopsis. Nucleic Acids Research, 35(SUPPL. 1), 4–6. 10.1093/nar/gkl783

Ordon, J., Gantner, J., Kemna, J., Schwalgun, L., Reschke, M., Streubel, J., Boch, J., & Stuttmann, J. (2017). Generation of chromosomal deletions in dicotyledonous plants employing a user-friendly genome editing toolkit. Plant Journal, 89(1), 155–168. 10.1111/tpj.13319

Palaniswamy, S. K., James, S., Sun, H., Lamb, R. S., Davuluri, R. V., & Grotewold, E. (2006). AGRIS and AtRegNet. A platform to link cis-regulatory elements and transcription factors into regulatory networks. Plant Physiology, 140(3), 818–829. 10.1104/pp.105.072280

Púčiková, V., Rohn, S., & Hanschen, F. S. (2023). Glucosinolate Accumulation and Hydrolysis in Leafy Brassica Vegetables Are Influenced by Leaf Age. Journal of Agricultural and Food Chemistry, 71(30), 11466–11475. 10.1021/acs.jafc.3c01997

Ritchie, M. E., Phipson, B., Wu, D., Hu, Y., Law, C. W., Shi, W., & Smyth, G. K. (2015). Limma powers differential expression analyses for RNA-sequencing and microarray studies. Nucleic Acids Research, 43(7), e47. 10.1093/nar/gkv007

Rodríguez-Leal, D., Lemmon, Z. H., Man, J., Bartlett, M. E., & Lippman, Z. B. (2017). Engineering Quantitative Trait Variation for Crop Improvement by Genome Editing. Cell, 171(2), 470–480.e8. 10.1016/j.cell.2017.08.030

Roodbarkelari, F., & Groot, E. P. (2017). Regulatory function of homeodomain-leucine zipper (HD-ZIP) family proteins during embryogenesis. In New Phytologist (Vol. 213, Issue 1, pp. 95–104). Blackwell Publishing Ltd. 10.1111/nph.14132

Rowe, H. C., Walley, J. W., Corwin, J., Chan, E. K. F., Dehesh, K., & Kliebenstein, D. J. (2010). Deficiencies in jasmonate-mediated plant defense reveal quantitative variation in Botrytis cinerea pathogenesis. PLoS Pathogens, 6(4), 1–18. 10.1371/journal.ppat.1000861

Schneider, C. A., Rasband, W. S., & Eliceiri, K. W. (2012). NIH Image to ImageJ: 25 years of image analysis. Nature Methods, 9(7), 671–675. 10.1038/nmeth.2089

Schu, H. (2003). Transcriptional control of nonfermentative metabolism in the yeast Saccharomyces cerevisiae. 139–160. 10.1007/s00294-003-0381-8

Sharma, R., Sharad, S., Minhas, G., Sharma, D. R., Bhatia, K., & Sharma, N. K. (2023). DNA, RNA isolation, primer designing, sequence submission, and phylogenetic analysis. In Basic Biotechniques for Bioprocess and Bioentrepreneurship. Elsevier Inc. 10.1016/B978-0-12-816109-8.00012-X

Shin, Y., Chane, A., Jung, M., & Lee, Y. (2021). Recent advances in understanding the roles of pectin as an active participant in plant signaling networks. Plants, 10(8), 1–22. 10.3390/plants10081712

Song, X., Ohbayashi, I., Sun, S., Wang, Q., Yang, Y., Lu, M., Liu, Y., Sawa, S., & Furutani, M. (2024). TCA cycle impairment leads to PIN2 internalization and degradation via reduced MAB4 level and ARA6 components in Arabidopsis roots. Frontiers in Plant Science, 15(December), 1–12. 10.3389/fpls.2024.1462235

Stuttmann, J., Barthel, K., Martin, P., Ordon, J., Erickson, J. L., Herr, R., Ferik, F., Kretschmer, C., Berner, T., Keilwagen, J., Marillonnet, S., & Bonas, U. (2021). Highly efficient multiplex editing: one-shot generation of 8× Nicotiana benthamiana and 12× Arabidopsis mutants. Plant Journal, 106(1), 8–22. 10.1111/tpj.15197

Tang, M., Li, B., Zhou, X., Bolt, T., Li, J. J., Cruz, N., Gaudinier, A., Ngo, R., Clark-Wiest, C., Kliebenstein, D. J., & Brady, S. M. (2021). A Genome-Scale TF-DNA Interaction Network of Transcriptional Regulation of Arabidopsis Primary and Specialized Metabolism. 10.1101/2021.05.13.443927

Tang, M., Li, B., Zhou, X., Bolt, T., Li, J. J., Cruz, N., Gaudinier, A., Ngo, R., Clark-Wiest, C., Kliebenstein, D. J., & Brady, S. M. (2021). A genome-scale TF–DNA interaction network of transcriptional regulation of Arabidopsis primary and specialized metabolism. Molecular Systems Biology, 17(11). 10.15252/msb.202110625

Wisecaver, J. H., Borowsky, A. T., Tzin, V., Jander, G., Kliebenstein, D. J., & Rokas, A. (2017). A global coexpression network approach for connecting genes to specialized metabolic pathways in plants. Plant Cell, 29(5), 944–959. 10.1105/tpc.17.00009

Yang, T. H. (2019). Transcription factor regulatory modules provide the molecular mechanisms for functional redundancy observed among transcription factors in yeast. BMC Bioinformatics, 20(Suppl 23), 1–16. 10.1186/s12859-019-3212-8

Yilmaz, A., Mejia-Guerra, M. K., Kurz, K., Liang, X., Welch, L., & Grotewold, E. (2011). AGRIS: The arabidopsis gene regulatory information server, an update. Nucleic Acids Research, 39(SUPPL. 1), 1118–1122. 10.1093/nar/gkq1120

Yu, H., Du, X., Zhang, F., Zhang, F., Hu, Y., Liu, S., Jiang, X., Wang, G., & Liu, D. (2012). A mutation in the E2 subunit of the mitochondrial pyruvate dehydrogenase complex in Arabidopsis reduces plant organ size and enhances the accumulation of amino acids and intermediate products of the TCA Cycle. Planta, 236(2), 387–399. 10.1007/s00425-012-1620-3

Zhang, W., Corwin, J. A., Copeland, D. H., Feusier, J., Eshbaugh, R., Cook, D. E., Atwell, S., & Kliebenstein, D. J. (2019). Plant–necrotroph co-transcriptome networks illuminate a metabolic battlefield. ELife, 8, 1–32. 10.7554/eLife.44279

Zhang, W., Corwinand, J. A., Copeland, D., Feusier, J., Eshbaugh, R., Chen, F., Atwell, S., & Kliebenstein, D. J. (2017). Plastic transcriptomes stabilize immunity to pathogen diversity: The jasmonic acid and salicylic acid networks within the Arabidopsis/Botrytis pathosystem open. Plant Cell, 29(11), 2727–2752. 10.1105/tpc.17.00348

